# Chemical genetics in *C. elegans* identifies anticancer mycotoxins chaetocin and chetomin as potent inducers of a nuclear metal homeostasis response

**DOI:** 10.1101/2024.02.15.579914

**Authors:** Elijah Abraham, A. M. Gihan K. Athapaththu, Kalina R. Atanasova, Qi-Yin Chen, Taylor J. Corcoran, Juan Piloto, Cheng-Wei Wu, Ranjala Ratnayake, Hendrik Luesch, Keith P. Choe

## Abstract

*C. elegans numr-1/2* (<u>nu</u>clear-localized <u>m</u>etal-responsive) is an identical gene pair encoding a nuclear protein previously shown to be activated by cadmium and disruption of the integrator RNA metabolism complex. We took a chemical genetic approach to further characterize regulation of this novel metal response by screening 41,716 compounds and extracts for *numr-1p::GFP* activation. The most potent activator was chaetocin, a fungal 3,6-epidithiodiketopiperazine (ETP) with promising anticancer activity. Chaetocin activates *numr-1/2* strongly in the alimentary canal but is distinct from metal exposure because it represses canonical cadmium-responsive metallothionine genes. Chaetocin has diverse targets in cancer cells including thioredoxin reductase, histone lysine methyltransferase, and acetyltransferase p300/CBP; further work is needed to identify the mechanism in *C. elegans* as genetic disruption and RNAi screening of homologs did not induce *numr-1/2* in the alimentary canal and chaetocin did not affect markers of integrator dysfunction. We demonstrate that disulfides in chaetocin and chetomin, a dimeric ETP analog, are required to induce *numr-1/2.* ETP monomer gliotoxin, despite possessing a disulfide linkage, had almost no effect on *numr-1/2*, suggesting a dimer requirement. Chetomin inhibits *C. elegans* growth at low micromolar levels and loss of *numr-1/2* increases sensitivity; *C. elegans* and Chaetomiaceae fungi inhabit similar environments raising the possibility that *numr-1/2* functions as a defense mechanism. There is no direct ortholog of *numr-1/2* in humans, but RNAseq suggests that chaetocin affects expression of cellular processes linked to stress response and metal homeostasis in colorectal cancer cells. Our results reveal interactions between metal response gene regulation and ETPs and identify a potential mechanism of resistance to this versatile class of preclinical compounds.

## INTRODUCTION

Metals are thought to bind half of all enzymes impacting nearly all biological functions. Heavy metal cadmium has no apparent physiological function but is a common environmental contaminant with broad molecular targets and severe consequences to development, behavior, immune function, and metabolism^1^. Understanding how animals respond to metals can improve treatment strategies, and small molecule modulators of metal homeostasis have the potential to therapeutically modify diverse biological processes^2^.

Cells respond to environmental stress by increasing transcription of cytoprotection genes. Genes induced rapidly during stress often contain few introns to bypass the need for pre-mRNA splicing^3^. In 2010, a pair of small intron-less genes named *<u>nu</u>clear localized <u>m</u>etal <u>r</u>esponsive-1/2* was discovered in *C. elegans* that have identical coding sequence and are highly induced by cadmium^4–5^. NUMR-1/2 protein sequence bears no resemblance to well-known metal responsive metallothionein metal ion chelators, protein chaperones, or previously studied ‘CaDmium Responsive’ (CDR) membrane proteins^4, 6–9^. Biochemical functions of NUMR-1/2 are unknown, but the protein was shown to localize to nuclei of intestine cells and to promote longevity and resistance to cadmium^4^. NUMR-1/2 has an RNA recognition motif at the N-terminus and serine and arginine repeats consistent with the serine/arginine-rich (SR) protein family that regulates RNA metabolism^6, 10^. NUMR-1/2 also has a histidine-rich region, raising the possibility of binding to metal ions and targeting to active sites of transcription^2, 11–12^.

To gain insights into how *numr-1/2* is regulated, we previously screened the *C. elegans* genome for dsRNA clones that activate a *numr-1* promoter reporter (*numr-1p::GFP*)^6^; the *numr-1* and *numr-2* promoters are 99% identical and drive reporter expression in the same tissues under that same conditons^4^. Genes identified by the screen were highly enriched for functions in RNA metabolism with silencing of integrator complex subunits causing the strongest *numr-1/2* activation^6^. Integrator is a metazoan-specific complex that functions in multiple steps of RNA metabolism including transcription termination and processing of non-coding small nuclear RNAs (snRNA)^13–15^. It was hypothesized that integrator disruption acts as a surveillance mechanism for environmental stress^6^.

To further characterize *numr-1/2* regulation and identify potential small molecule modulators of metal homeostasis, we screened 41,716 compounds and natural product extracts from six libraries for *numr-1p::GFP* activation in *C. elegans*; we identified 18 candidates and five were confirmed to be active. The mycotoxin chaetocin is a potent inducer of *numr-1/2* in the alimentary canal with low *in vivo* toxicity in *C. elegans*. Chaetocin is a homodimeric 3,6-epidithiodiketopiperazine (ETP) (Fig. S1) with diverse biological targets and promising anticancer activity^16–24^. Robust activation of *numr-1/2* provides an opportunity to gain new insights into chaetocin bioactivity and metal response regulation. Surprisingly, chaetocin suppresses canonical cadmium responsive metallothionein genes. Chaetocin does not affect expression of markers for integrator complex dysfunction, and genetic loss of worm homologs of previously identified chaetocin targets thioredoxin reductase and worm p300/CBP orthologue does not mimic effects of the compound on *numr-1/2*. Heterodimeric ETP chetomin has similar activity as chaetocin, but not monomeric ETP gliotoxin nor disulfide depleted analogs (Fig. S1) suggesting that the disulfide ETP warhead is required, but not sufficient, to induce *numr-1/2*. Chetomin inhibited *C. elegans* growth at low micromolar concentrations and loss of *numr-1/2* increased sensitivity. There is no direct human ortholog of NUMR-1/2, but we used RNAseq of a colorectal cell line to explore the possibility that chaetocin and chetomin effects on metal-homeostasis gene expression are conserved; processes enriched among the genes affected were immune responses and hemostasis, which are tightly linked to zinc homeostasis. Taken together, our results reveal interactions between dimeric ETPs and metal response gene expression and identify *numr-1/2* as a nematode response to ETPs that promotes resistance.

## MATERIALS AND METHODS

### C. elegans strains and culture conditions

Unless stated otherwise, all *C. elegans* strains were maintained at 20°C on nematode growth medium (NGM) agar plates with OP50 bacteria using standard methods^25^. Strains used in the study include: N2 Bristol, QV320 *qvIs4[numr-1p::GFP];vsIs33[dop-3p::RFP]*, QV325 *dpy-5 (e61);qvIs4;vsIs33,* QV335 *numr-1;numr-2(sybDf10)*, VZ21 *trxr-2(ok2267)* III; *trxr-1(sv47)* IV, QV380 *qvIs15[pCL26 mtl-2p::GFP*] (outcrossed from CL2122), and QV381 *numr-1;numr-2(sybDf10);qvIs15*.

A translational reporter for NUMR-1/2 was created by injecting N2 with a PCR fragment containing 454 bp of primer and the entire coding region of *numr-1/2* fused to mCherry (QV315 *zjEx141[numr-1/2p::numr-1/2::mCherry; myo-2p::tdTomato]*); mCherry was used to allow future colocalization with GFP reporters.

Rescue and control lines were generated by injecting N2 and *numr-1/2* worms with 15 ng/µl of *numr-1* and *numr-2* gDNA PCR fragments that include the entire upstream promoter, coding region, and 454 or 486 bp of downstream sequence, respectively (QV376 kbEx150[*numr-1, numr-2* gDNA; *myo-2p::tdTomato*], QV377 kbEx151[*numr-1, numr-2* gDNA; *myo-2p::tdTomato*], QV378 *numr-1;numr-2(sybDf10);*kbEx152[*numr-1, numr-2* gDNA; *myo-2p::tdTomato*], and QV379 *numr-1;numr-2(sybDf10);*kbEx153[*numr-1, numr-2* gDNA; *myo-2p::tdTomato*]).

### Fluorescence screening

Screening was conducted for *in vivo* activation of *numr-1p::GFP* in *C. elegans* strain QV325 similar to our prior screens with some modifications^26–27^. Compounds were dispensed with a JANUS liquid handling system (Perkin Elmer) at 75 μl into black 96-well (commercial and University of Florida proprietary libraries) or at 30 μl into 384-well plates (NCI Natural Product Discovery library) in liquid NGM buffer containing either 0.01% Tween-20 or Triton X-100 to limit worm adherence to plastic. Worms were synchronized by standard bleach treatment^28^ and fed NA22 on peptone-rich agar plates until the L4 to young adult stage. *In vivo* induction of *numr-1p::GFP* in *C. elegans* requires the presence of bacteria food to promote ingestion. Concentrated OP50 bacteria was prepared as we have described previously^26^, washed in NGM buffer with either 0.01% Tween-20 or Triton X-100, and heat-shocked in a water bath to 56°C for 5 min. Heat-treated bacteria (final OD_600_ 0.8-1.25) and 65-75 worms in 25 μl were added to compounds in each well of 96-well plates or 40-50 worms in 15 μl for each well of 384-well plates. Final concentrations of compounds were 1 to 50 μM with a maximum DMSO concentration of 1% (Table S1). Plates were sealed with breathable tape and incubated for 16-24 hours while rocking at 20°C. The presence of bacteria and localized tissue-specific *numr-1p::GFP* made plate reader quantification of fluorescence unreliable. Therefore, we screened for fluorescence induction manually with a Zeiss Stemi SV12 microscope. Fluorescence of *numr-1p::GFP* is nearly absent under basal conditions at low magnification when viewed manually; compounds that caused visible activation of fluorescence in at least 50% of worms were scored as hits. Structures of confirmed *numr-1/2* inducers are in Figure S1.

### C. elegans imaging and growth assays

*C. elegans* fluorescent reporter strains were exposed to compounds in NGM buffer as described for screening using clear 96-well plates except they were incubated with lids in humified cambers. After incubation, worms were mounted on slides with 1.5-2% agarose pads using hair picks in 5-25 mM tetramizole or levamisole as anesthesia. Worms were imaged with an Olympus BX60 microscope with a Zeiss AxioCam MRm camera. For *numr-1p::GFP*, whole worms were imaged at 10x magnifications and average pixel intensity of induced tissues was quantified using ImageJ^29^. For *mtl-2p::GFP*, average fluorescence of the whole intestine was quantified at 4x magnification.

For growth assays, wildtype N2 or QV335 *numr-1/2* mutant worms were bleach synchronized and added to NGM buffer with OP50 (final OD_600_ 0.8-1.25) in 96-well plates at the L1 stage. Two days later, relative length of worms was measured with an LP Sampler and COPAS BioSort and averaged for each well. In a control experiment, cell division and metabolic activity of OP50 were suppressed by treatment with 0.25% paraformaldehyde for 1 h at 37°C and washed as described previously^30–31^. For *numr-1/2* genetic rescue experiments, fluorescence of co-injection marker *myo-2p::tdTomato* was used to identify worms with arrays.

### *RT-qPCR* measurements of *numr-1/2* and representative stress-response genes

For RT-qPCR, wildtype N2 or mutant worms without transgene reporters were exposed to compounds in 96-well plates as described above for imaging. Because stress response gene expression can change quickly and may be sensitive to processing, we collected worms by picking into lysis buffer and freezing as we and others have described previously with some modifications^32–35^; this approach is rapid, selective for developmental stage and condition, and has few processing steps limiting artifacts that can be introduced during bulk collection and washing of entire plates of worms. At the end of exposures, worms and incubation buffer containing compounds or vehicle controls were transferred to the surface of agar plates by pipet; for each replicate, five or ten young adult worms were picked to tubes containing lysis buffer and proteinase K^32^ and frozen by placement in a pre-chilled rack in an ultracold freezer. Worms were lysed at 65°C for 11 min and proteinase K according to manufacturer recommendations. After lysis, genomic DNA was digested with DNase (Thermo Fisher AM1907 or EN007) and cDNA was synthesized with GoScript Reverse Transcriptase (Promega A5001) according to manufacturer protocols. qPCR assays were run with Forget-Me-Not Master Mix (Biotium 31044) in an Eppendorf Realplex 2 or BioRad CFX96 machine. Gene expression was normalized to housekeeping gene *rpl-2* and to a value of 1.0 for controls with the delta-delta Ct method. Primers are listed in Table S2.

### RNAi experiments

Worms were fed *Escherichia coli* [HT115(dE3)] engineered to produce double-stranded RNA (dsRNA) homologous to target *C. elegans* genes on agar plates as described previously^36^. The control RNAi clone was pPD129.36 (L4440) and it encodes a 202-bp dsRNA that is not homologous to any *C. elegans* gene. The *cbp-1* dsRNA clone was obtained from the MRC library and verified by sequencing. One set of synchronized L1 larvae were fed dsRNA-expressing bacteria until the L4 stage (before obvious organogenesis defects) and imaged as described above. A second set of L1s were fed control dsRNA-expressing bacteria until the L3 stage then collected in NGM buffer and divided between plates with either control or *cbp-1* dsRNA-expressing bacteria; these worms were imaged on the first and third days of adulthood as described above.

### Human cell whole-transcriptome RNA sequencing

HCT116 cells were seeded in 6-well plates at 2 × 10^5^ cells/well in Dulbecco’s Modified Eagle Medium supplemented with 10% fetal bovine serum and 1% antibiotic-antimycotic solution. After attaching and acclimating overnight, cells were treated with vehicle control (DMSO, 0.5%), chaetocin, chetomin, or respective S-methyl products at 500 nM for 5 or 12 h. Total RNA was extracted using an RNeasy Mini Kit (Qiagen 74104) and sent to the University of Florida Interdisciplinary Center for Biotechnology Research (UF ICBR) for mRNA isolation, library construction (New England Biolabs, E7490 and E7760), and sequencing of paired 150 bp reads in an Illumina NovaSeq. Quality control was conducted with Cutadapt, reads were mapped to genome version GRCh38 with STAR, counts were processed using HTSeq and Samtools to remove PCR duplicates and count unique reads, and differential expression was analyzed with DESeq2. Functional enrichment analysis of co-regulated genes was conducted with DAVID^37^. Raw sequence data are available at GEO GSE232639.

### Statistical analyses

RT-qPCR and fluorescence data were analyzed with Students *t-tests* and P-values and were corrected for multiple comparisons with Benjamini-Hochberg adjustments. Dose response curves were generated with a non-linear log(dose) variable slope model in GraphPad Prism 5.04. Underlying numeric data are available at figshare https://doi.org/10.6084/m9.figshare.25207175.v1.

## RESULTS

### In vivo fluorescence screening identified inducers of numr-1p::GFP

A scheme of the screening approach and the structure of the most potent *numr-1/2* inducer identified, chaetocin, is shown in Fig. 1. Synchronized transgenic *numr-1p::GFP* reporter worms were collected from agar plates at the L4 to young adult stage, washed, and added to black 96 or 384-well plates with compounds in liquid NGM media. To promote ingestion, the standard lab diet *E. coli* strain OP50 was also included after exposure to a five-minute water bath heat-shock at 56°C to reduce biochemical activity^38^. Although mRNA of stress response genes like *numr-1/2* are induced within a few hours of exposure to environmental stimuli, accumulation of fluorescent reporter proteins is delayed by several hours as they require translation and maturation of GFP^4, 36, 39–43^. Accordingly, we incubated reporter worms with compounds for 16-24 h before analysis; a constitutive *dop-3p::RFP* reporter was included to allow focusing on worms within black plates. Fluorescence was scored manually because the presence of bacteria and tissue-specific *numr-1p::GFP* induction made plate reader quantification unreliable. When viewed with a dissecting microscope, basal *numr-1p::GFP* fluorescence is nearly undetectable; compounds were scored as hits if they caused obvious increases in *numr-1p::GFP* fluorescence in at least half of the worms within a well.

**Figure 1.**
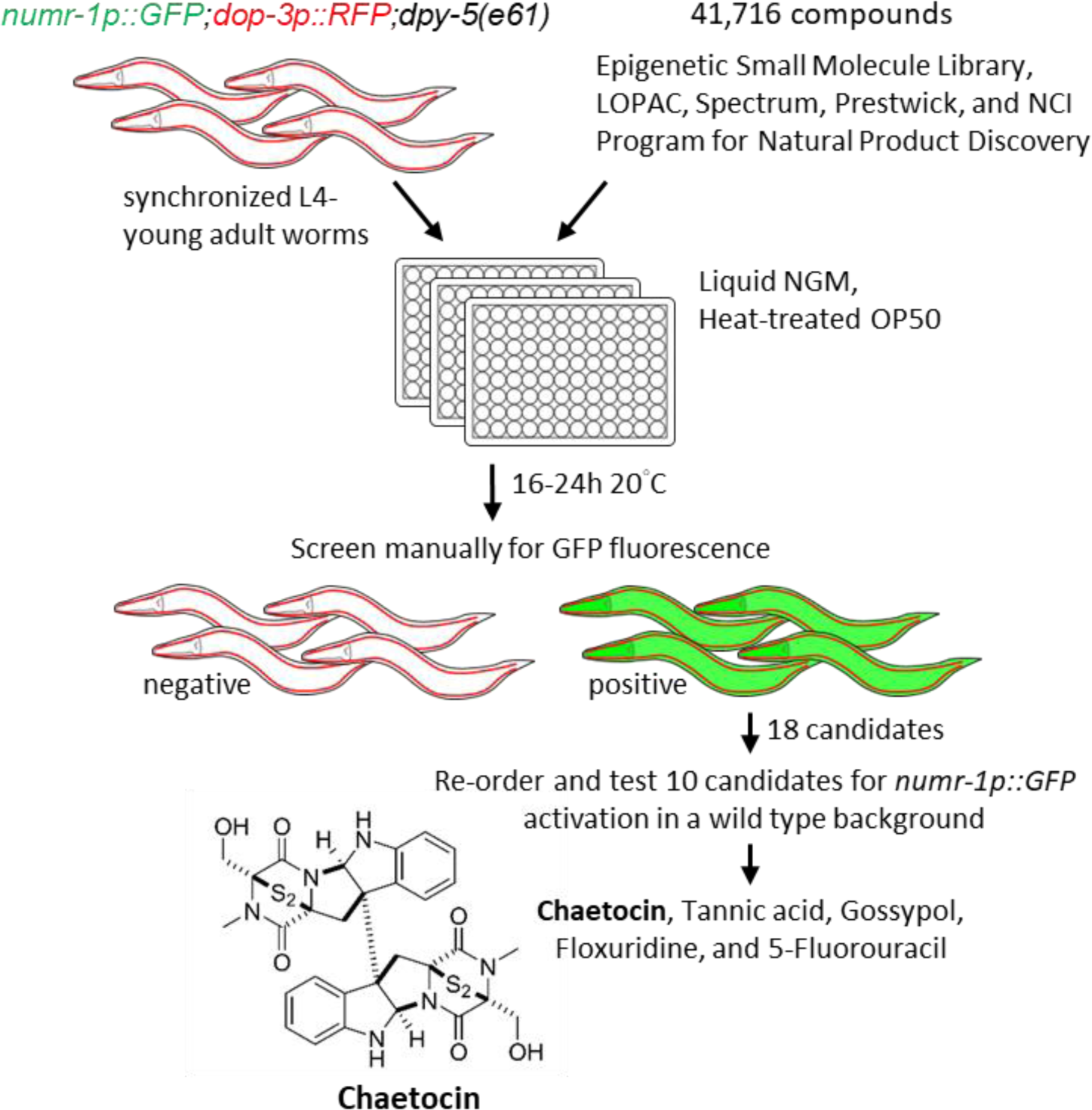
Scheme of initial screening for *numr-1p::GFP* inducers. Initial screening for *numr-1p::GFP* activation was conducted manually in worms with a *dpy-5* cuticle collagen gene mutation and a constitutive RFP reporter. Ten candidates were reordered and tested and five induced *numr-1p::GFP* in a wild type background without acute toxicity. Chaetocin was characterized further.

We identified 18 distinct candidate small molecules by screening 41,716 compounds or natural product fractions from six libraries for *numr-1p::GFP* activation; libraries were Epigenetic Small Molecule Library (Cayman Chemicals), LOPAC (Sigma), Spectrum (Microsource Discovery Systems, Inc.), Prestwick Chemical Library, University of Florida proprietary internal natural and synthetic compound libraries, and a portion of the NCI Program for Natural Product Discovery (NPNPD) pre-fractionated library (Table S1). Absorption of many small molecules into *C. elegans* is restricted by a collagenous barrier cuticle^44^; our initial screening was conducted with high compound doses (50 μM), heat-treated bacteria to promote ingestion, and a *dpy-5(e61)* cuticle collagen mutation that enhances permeability^40^. Even in these conditions, poor penetration may have contributed to a low hit rate of 0.04% and it is possible that we missed some compounds with the potential to induce *numr-1p::GFP*. We reordered and tested 10 candidates in a wild type background, which is the standard laboratory genetic background and is most convenient for further characterization; six induced *numr-1p::GFP*, but one (actinomycin D) was highly toxic and not characterized further (Figs. 1, 2A, S1, and Table S3). Fungal product chaetocin and plant polyphenols tannic acid and gossypol activated *numr-1p::GFP* in the pharynx and/or intestine (Figs. 2 and S2) matching the location of *numr-1/2* induction by metals^4, 6^. Floxuridine and 5-fluorouracil are well-characterized inhibitors of DNA synthesis^45–47^ and they only activated *numr-1p::GFP* in late embryos (Figs. 2 and S2); embryos are normally sites of rapid DNA and RNA synthesis and were arrested in the presence of these DNA synthesis inhibitors.

**Figure 2.**
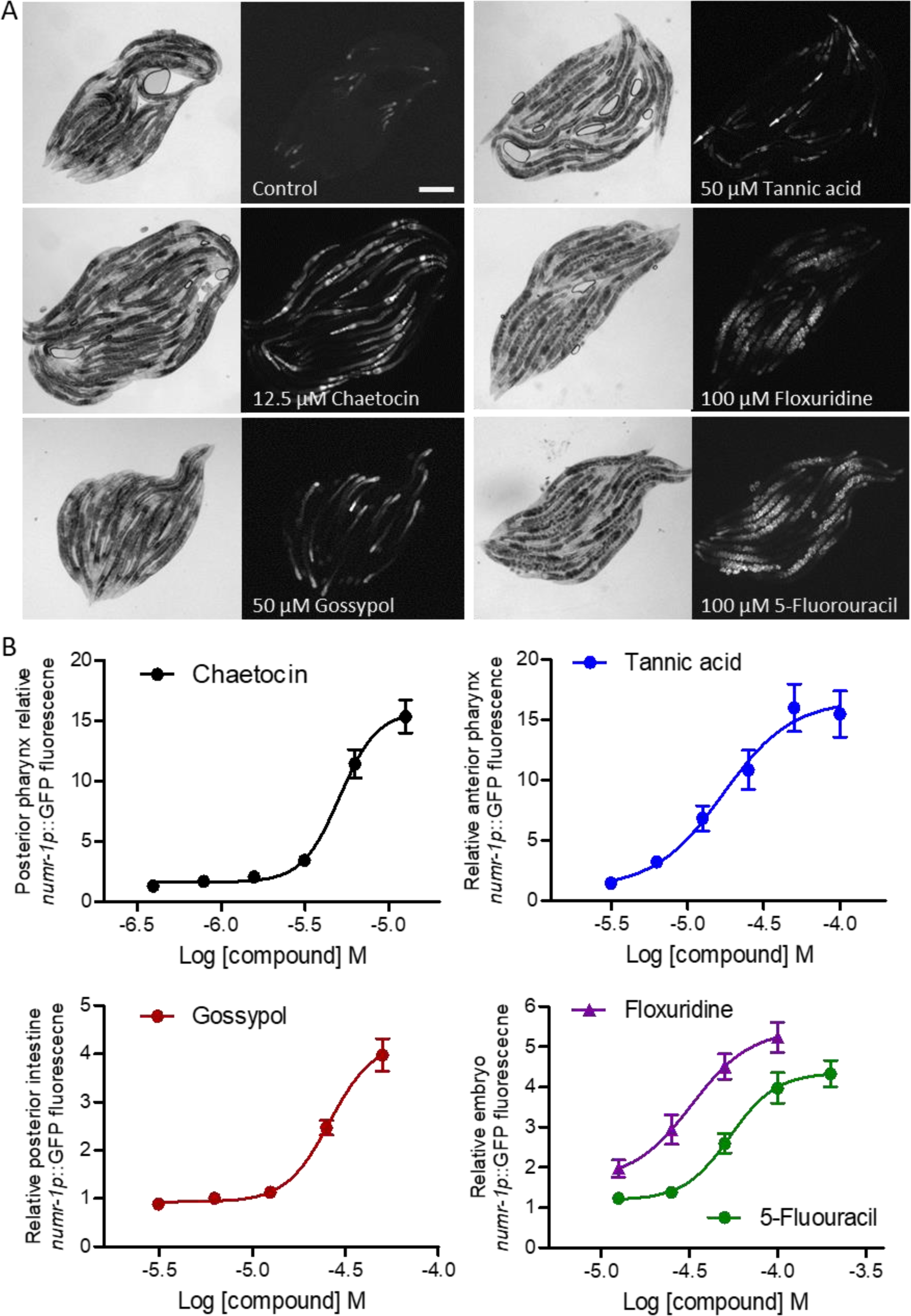
Five confirmed *numr-1p::GFP* inducers were tested at multiple doses in a wildtype background. (A) Representative bright field and fluorescence micrographs of *numr-1p::GFP* worms exposed to active compounds. Scale bar is 200 µm. (B) Dose-responses for relative *numr-1p::GFP* fluorescence in tissues most strongly induced for each compound. Mean ± standard error normalized to 1.0 for vehicle controls. N = 25-36 individual worms from two trials at each concentration.

We next tested *numr-1p::GFP* inducers at multiple doses. After incubation for 18-24 h, worms were mounted on slides, imaged, and GFP fluorescence was quantified in individual worms in tissues with the most robust and consistent induction as follows: posterior pharynx bulb for chaetocin, anterior pharynx for tannic acid, posterior intestine for gossypol, and embryos for floxuridine and 5-fluorouracil. Although low-throughput and labor intensive, we used this approach because *numr-1p::GFP* is tissue-specific and variable between compounds (Figs. 2A and S2) making results from plate reading of worm populations or scans of whole worms with a BIOSORT flow cytometer insensitive and potentially misleading. All five compounds activated *numr-1p::GFP* dose-dependently with the following estimates for EC_50_: chaetocin (5.1 µM), tannic acid (17.1 µM), gossypol (26.0 µM), floxuridine (32.5 µM), and 5-fluorouracil (53.7 µM) (Fig. 2B); the highest doses shown are those that were tolerated well by worms and without visible precipitation. Higher doses of tannic acid were toxic and reduced activity, and higher doses of floxuridine and 5-fluorouracil blocked embryo formation; higher doses of chaetocin and gossypol were not fully soluble. Variation in *numr-1p::GFP* fluorescence levels among individual worms is shown in a scatter plot in Figure 3A for each compound at near maximal activation doses; a logarithmic scale is used because of the large range in means between compounds. All means were significantly greater than a normalized value of 1.0 for within-trial vehicle controls.

**Figure 3.**
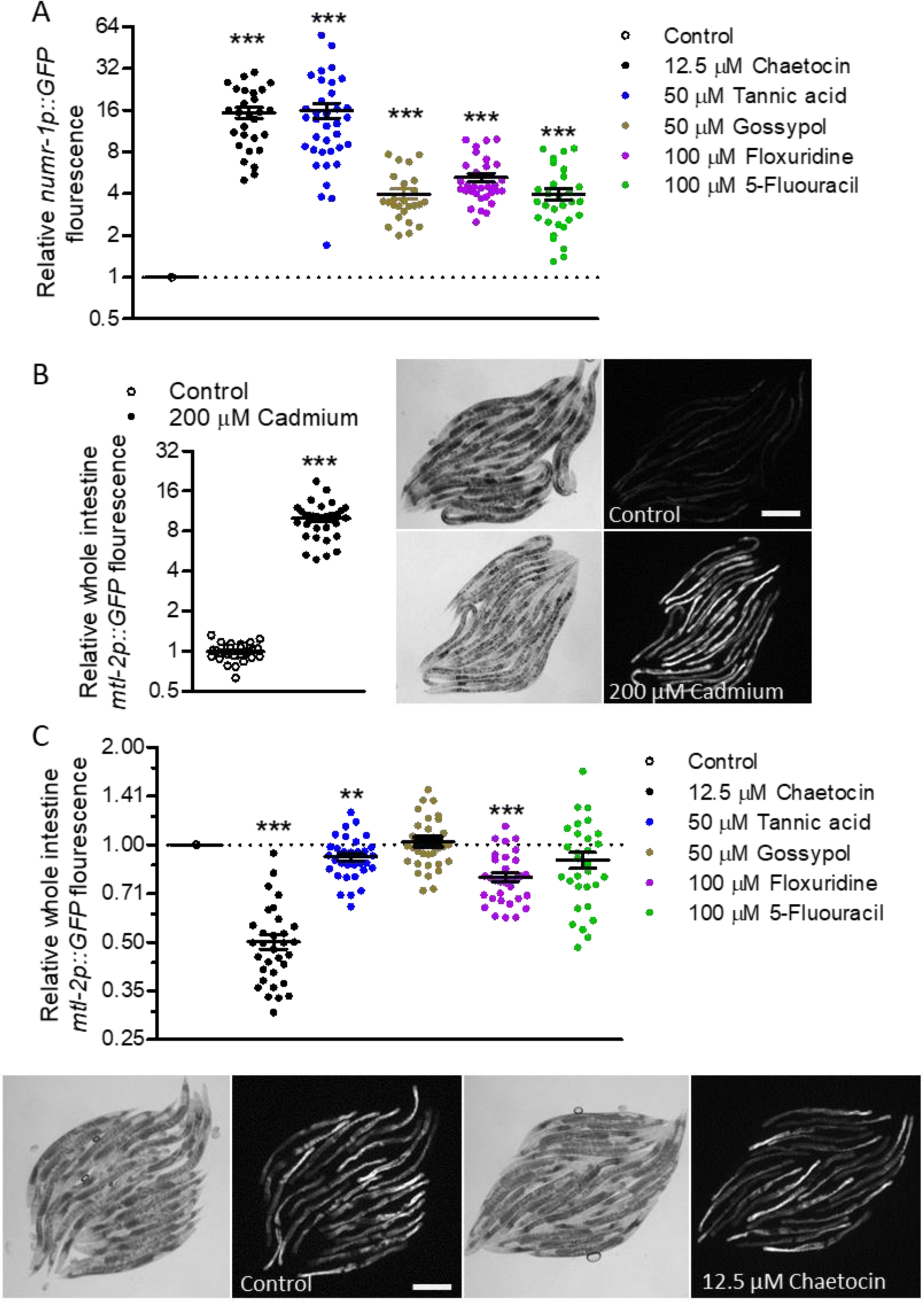
The five *numr-1p::GFP* inducers do not activate *mtl-2p::GFP*. (A) Scatter plot of relative *numr-1p::GFP* fluorescence in individual worms at near-maximal activation concentrations; because some compounds were tested in different trials and activated fluorescence in different tissues, all means were compared to a normalized within-trial value of 1.0 for vehicle controls. (B) Scatter plot of relative *mtl-2p::GFP* fluorescence in individual worms with 200 µM cadmium and representative bright field and fluorescence micrographs of *mtl-2p::GFP* worms. Scale bar is 200 µm. (C) Scatter plot of relative *mtl-2p::GFP* fluorescence in individual worms with the same compound concentrations as in panel A and representative bright field and fluorescence micrographs of *mtl-2p::GFP* worms exposed to chaetocin; all means were compared to a normalized within-trial value of 1.0 for vehicle controls. Note that longer camera exposure times are shown in C than B. (A-C) Mean ± standard error. \*\**p* < 0.01 and \*\*\**p* < 0.001 versus normalized vehicle control values of 1.0. N = 25-36 individual worms from two trials for each condition.

### A canonical metallothionine gene promoter is not activated by *numr-1p::GFP* inducers

Metallothioneins are small cysteine-rich proteins that regulate cell homeostasis by reversibly binding metal ions^5, 48^. In *C. elegans*, metallothionein gene *mtl-2* is expressed throughout the intestine and is activated strongly by high levels of metals^9, 42^. As expected and shown in Figure 3B, a previously characterized GFP transgene driven by the *mtl-2* promoter (*mtl-2p::GFP*)^49–50^ is highly induced by cadmium. To determine if *numr-1/2* inducers also activate this metallothionein gene promoter, we measured *mtl-2p::GFP* fluorescence after exposure to the same concentrations used in Figure 3A. As shown in Figure 3C, none of the *numr-1/2* inducers increased *mtl-2p::GFP* fluorescence; alternatively, chaetocin, tannic acid, and floxuridine reduced *mtl-2p::GFP* fluorescence by 50, 8, and 21%, respectively; note that longer camera exposure times are shown in Fig. 3C than 3B. These results suggest that these compounds induce *numr-1/2* without also activating the canonical metallothionein metal response.

### Chaetocin and tannic acid induce endogenous numr-1/2 mRNA

Cadmium induces *numr-1/2*, but also activates many other stress responses consistent with broad bioactivity^4–6, 48, 51^. A prior microarray study reported that chronic growth in 100 µM tannic acid increases *numr-1/2* expression by 75-fold^52^, but expression data is not available for shorter exposures or for chaetocin at any dose. To verify upregulation of endogenous mRNA by the two most robust inducers (chaetocin and tannic acid) and assess specificity among stress responses, we measured relative mRNA levels of *numr-1/2* and genes representative of core metal (*mtl-1* and *mtl-2*), endoplasmic reticulum unfolded protein (*hsp-4*), and detoxification (*gst-4*) responses *via* RT-qPCR. As described above, stress response genes like *numr-1/2* are activated strongly within a few hours of exposure to stimuli, several hours before fluorescent protein reporters accumulate^4, 6, 42^; therefore, similar to prior studies with 4 h of cadmium^42^, worms were treated for 5 h to capture early changes in gene expression while reducing long-term indirect effects; the doses were the same as in Figure 3. Five-hour exposure to 12.5 µM chaetocin or 50 µM tannic acid upregulated endogenous *numr-1/2* mRNA over 50-fold without activating the other stress response genes measured (Figs. S3A-B).

Consistent with the reporter results in Figure 3C, neither compound induced *mtl-2* mRNA; both compounds suppressed *mtl-2* mRNA and chaetocin also suppressed *mtl-1* (Fig. S3A-B). Therefore, chaetocin and tannic acid can strongly induce *numr-1/2* without activating metallothionein genes. Tannic acid is a well-studied plant polyphenol and traditional medicine with diverse bioactivities including increased longevity in *C. elegans*^52–55^; tannic acid and other polyphenols can form complexes with metal ions providing a potential mechanism for changes to *numr-1/2* regulation^55^.

Chaetocin is a 3,6-epidithiodiketopiperazine (ETP) first isolated as a secondary metabolite of fungus *Chaetomium minutum* in 1970^56^; it has promising activity against cancer cells but mechanisms of action are likely diverse and not fully understood^16–18, 23, 57^. We focus the remainder of this study on chaetocin and analogs. Fluorescent fusion protein reporters for NUMR-1/2 are localized to intestine nuclei when induced by cadmium^4^; the intestine has the largest somatic cells and nuclei in *C. elegans* as a result of endoreduplication making nuclear localization of protein straightforward to visualize even at low magnification^58^. As shown in Figure S3C, a NUMR-1/2::mCherry fusion protein localizes to intestine nuclei when induced by chaetocin.

### Interactions between chaetocin and cadmium

We next tested interactions between chaetocin and cadmium. Cadmium activates *numr-1/2* and some previously characterized activities of ETP natural products are hypothesized to involve displacement of physiological zinc^59–61^. At 200 μM, metal chelator TPEN strongly reduces free cytosolic zinc levels and disrupts *C. elegans* reproduction^62–63^. As shown in Figure 4A, 200 μM TPEN increased *numr-1p::GFP* fluorescence by 1.3-fold; this is statistically significant, but far less than the over 15-fold induction by 12.5 μM chaetocin (Fig. 3A) suggesting that *numr-1/2* is not strongly activated by broad depletion of cellular zinc.

**Figure 4.**
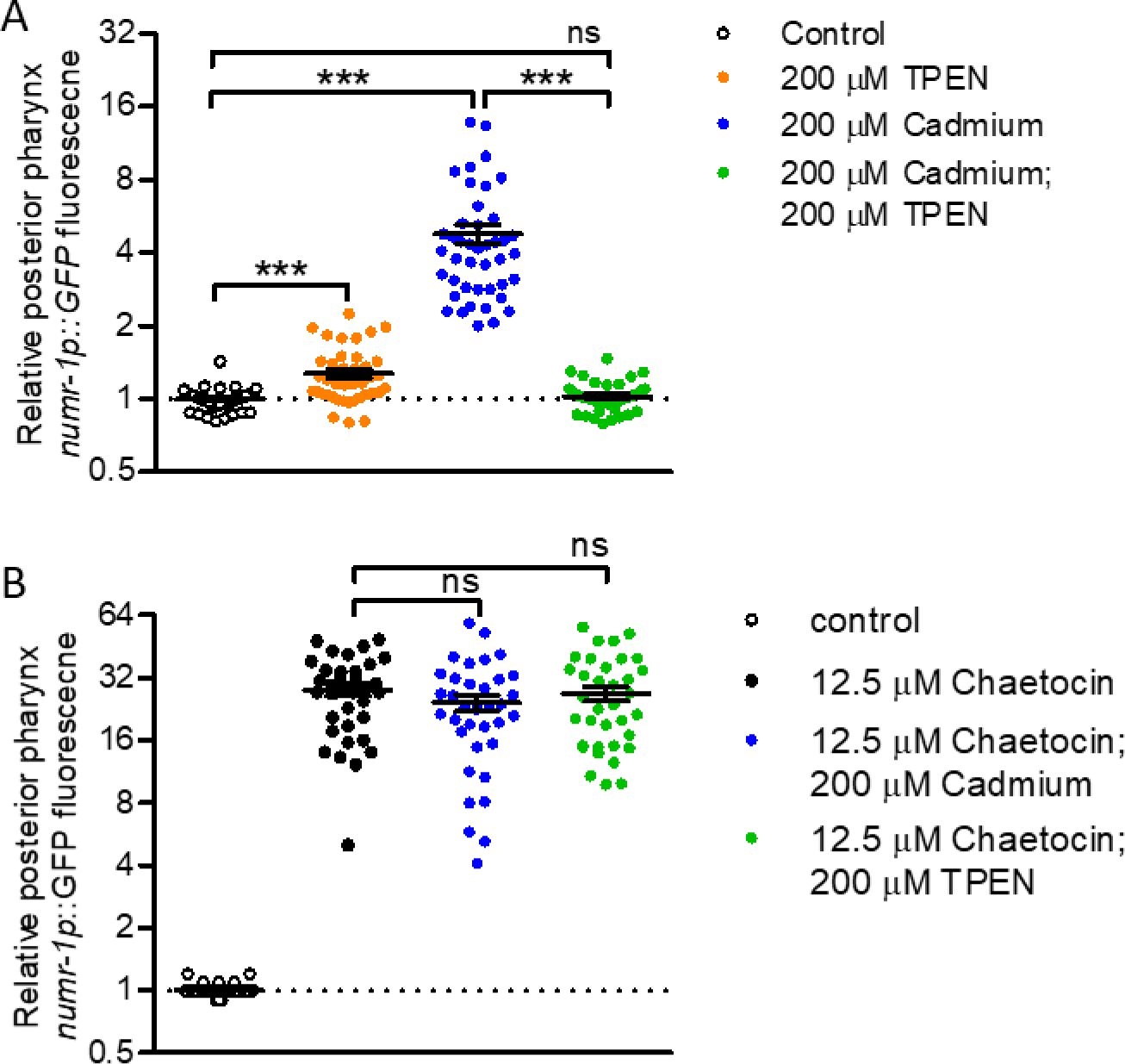
Interactions between chaetocin, cadmium, and metal chelator TPEN. (A) Relative *numr-1p::GFP* fluorescence in worms exposed to cadmium, TPEN, or both. (B) Relative *numr-1p::GFP* fluorescence in worms exposed to chaetocin alone or with cadmium or TPEN; all means were significantly different from vehicle controls (P < 0.001). (A-B) Mean ± standard error, \*\*\**p* < 0.001 and ns, not significant. N = 32-43 individual worms from two trials.

As expected, 200 μM cadmium induced *numr-1p::GFP* in the posterior pharynx bulb over 4-fold; 200 μM TPEN blocked *numr-1p::GFP* activation by 200 μM cadmium consistent with transition metal ion chelation (Fig. 4A); alternatively, 200 μM TPEN had no effect in the presence of 12.5 μM chaetocin (Fig. 4B). As shown in Figure 4B, 200 μM cadmium was not additive with 12.5 μM chaetocin raising the possibility that chaetocin and cadmium may share overlapping mechanisms for regulation of *numr-1/2*.

### Markers of integrator dysfunction are not induced by chaetocin

We previously demonstrated that *numr-1/2* expression is highly sensitive to disruption of integrator function^6^. Loss of integrator genes was previously demonstrated to reduce snRNA processing and dramatically increase unconventional transcription of RNA chimeras of snRNA and downstream gene loci^15^; increases in these ‘read-through’ transcripts provide a sensitive marker for integrator dysfunction^15^. As expected, we observed over 50-fold increases in RNA levels for three of these read-through transcripts when integrator subunit 4 (*ints-4*) was silenced (Fig. S4A). Alternatively, 12.5 µM chaetocin for 5 h had no effect on any of these read-through transcripts (Fig. S4B).

### Loss of thioredoxin reductases or p300/CBP-1 does not mimic chaetocin

Previous work identified chaetocin as a thioredoxin reductase inhibitor, and it was posited that this activity induced cell death by promoting reactive oxidative species in cancer cells^16, 19^. *C. elegans* has two thioredoxin reductases, TRXR-1 and TRXR-2; *trxr-1(sv47);trxr-2(ok2267)* double deletion mutants are superficially wildtype^64^. Loss of both thioredoxin reductases had no effect on *numr-1/2* expression and increased *mtl-2* slightly (Fig. S4C).

Chaetocin was previously shown to inhibit transcription factor binding by histone acetyltransferase co-regulator p300 in colorectal and hepatocellular carcinoma cells^59^. *C. elegans* has a single ortholog of p300/CBP named CBP-1^65^. Loss of *cbp-1* is pleiotropic causing gonadogenesis defects, sterility, and embryonic lethality and cannot be maintained as a homozygote. We instead used RNAi to silence *cbp-1.* The effects of *cbp-1* loss on organ development make interpretation of whole worm mRNA levels problematic; we instead compared *numr-1p::GFP* fluorescence in the pharynx. One set of worms were fed *cbp-1* dsRNA at the first larval stage (L1) and were imaged at the last larval stage (L4) before gross organ defects became obvious; another set of worms were fed *cbp-1* dsRNA at the third larval stage (L3) and were imaged on the first day of adulthood. As shown in Figure S4D, *cbp-1* RNAi did not induce pharynx *numr-1p::GFP* in either set of worms; there was also no induction of *numr-1p::GFP* in the intestine (Figs. S4E-F). Worms treated with *cbp-1* RNAi from L3 to day 3 of adulthood developed disorganized gonads as expected; *numr-1p::GFP* fluorescence was induced in these worms, but it was localized to amorphous cells outside of the pharynx and intestine (Fig. S4G).

Chaetocin inhibits histone lysine methyltransferases SUV39H1 and G9a^22–23, 66^. If chaetocin affects metal responsive genes by this mechanism, then loss of a target methyltransferase should have similar effects. The *C. elegans* genome is predicted to encode over 40 methyltransferases^67^; 24 of these are included in our RNAi library, but none activated *numr-1p::GFP* in our previous genome-wide screen^6^. We re-screened these methyltransferase dsRNA clones in isolation (Table S4), but again failed to identify any that activate *numr-1/2p::GFP*. Further work including unbiased suppressor screens are needed to define the mechanism of *numr-1/2* regulation by chaetocin; we focus the rest of this study on testing chemical analogs and the function of *numr-1/2*.

### Heterodimer chaetocin analog chetomin induces numr-1/2

To gain insights on how the structure of chaetocin contributes to *numr-1/2* induction, we first conducted dose responses with related compounds that are commercially available (Fig. 5A). Monomeric ETP gliotoxin was previously shown to affect behavior and aging in *C. elegans*^68–69^, but had very little effect on pharynx *numr-1p::GFP* compared to chaetocin (1.3 and 21-fold, respectively at 12.5 μM, Fig. 5B) indicating that a disulfide functionality is not sufficient for robust induction. Alternatively, heterodimeric ETP chetomin induced *numr-1p::GFP* in the anterior intestine (Figs. 5C-D and S5A) with an estimated EC_50_ of 1.4 μM, but was toxic to worms above 10 µM limiting maximal total fluorescence. Gliotoxin also had minimal effect on *numr-1p::GFP* in the intestine (Fig. 5C). With RT-qPCR, 3.15 μM chetomin increased endogenous *numr-1/2* mRNA and suppressed *mtl-1* and *mtl-2 mRNA* (Fig. S5B).

**Figure 5.**
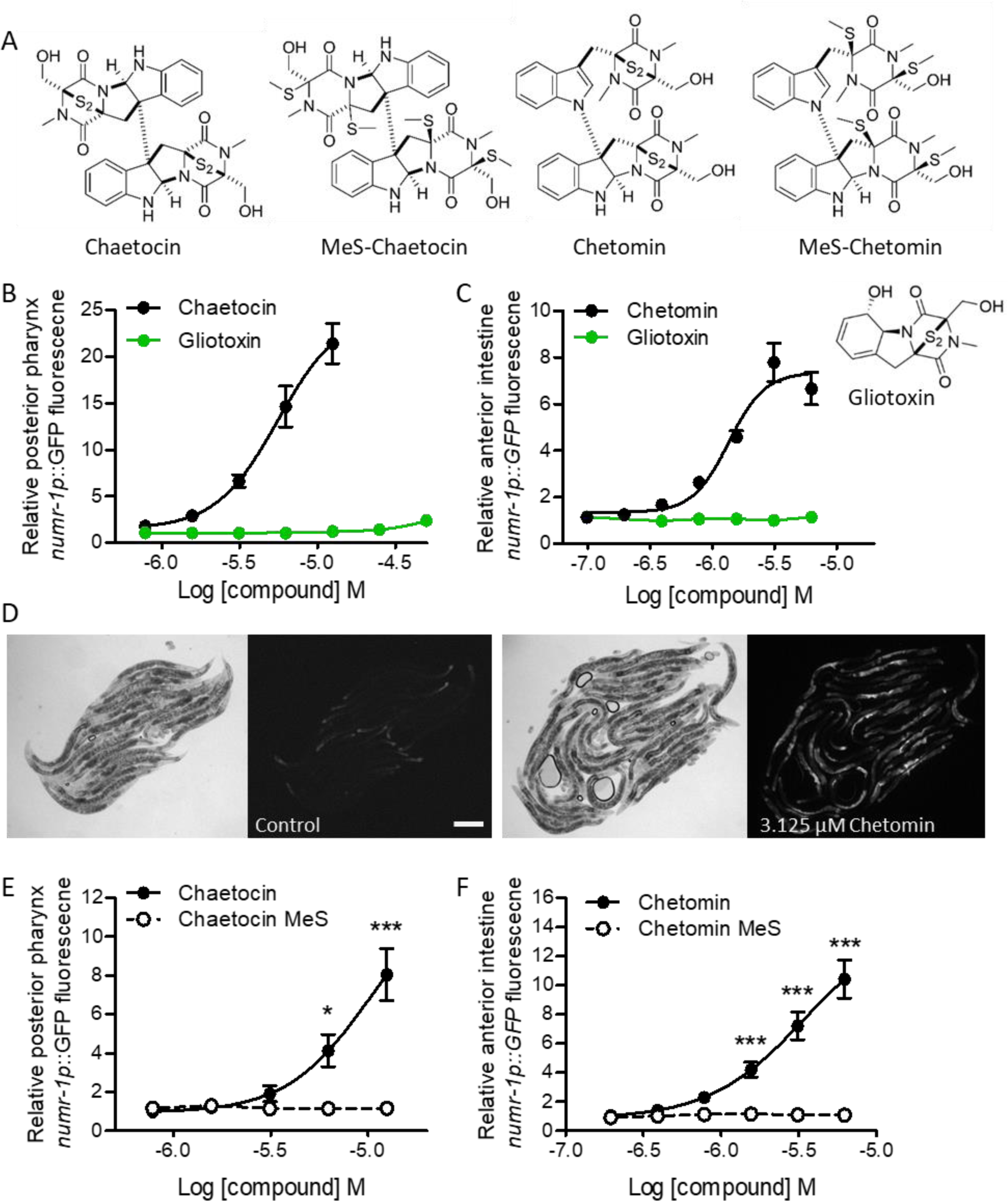
Heterodimer chaetocin analog chetomin induce *numr-1p::GFP* in the intestine. (A) Structures for chaetocin, chetomin, gliotoxin, and methylated sulfur analogs MeS-chaetocin and MeS-chetomin. (B) Relative *numr-1p::GFP* fluorescence for chaetocin and gliotoxin in the posterior pharynx. Mean ± standard error. N = 22-24 worms from two trials for each compound at each dose. (C) Relative *numr-1p::GFP* fluorescence for chetomin and gliotoxin in the anterior intestine. Mean ± standard error. N = 21-32 worms per dose from one trial for chetomin and two trials for gliotoxin. (D) Representative bright field and fluorescence micrographs of *numr-1p::GFP* worms exposed to chetomin. Scale bar is 200 µm. (E) Relative *numr-1p::GFP* fluorescence for chaetocin and MeS-chaetocin in the posterior pharynx bulb. N = 9-10 worms from 1 trial per dose of each compound. (F) Relative *numr-1p::GFP* fluorescence for chetomin and MeS-chetomin in the anterior intestine. N = 8-12 worms from 1 trial per dose of each compound. \**p* < 0.05 and \*\*\**p* < 0.001 versus MeS-chaetocin or MeS-chetomin.

Structure-activity relationships have demonstrated that disulfide-deficient analogs of chaetocin are not active against G9a^70–71^. We next tested if disulfide bridges of chaetocin and chetomin ETP cores are required for *numr-1/2* induction; disulfides were removed by methylation as described in Supplementary Information and Table S5. As shown in Figures 5E-F, methylated sulfide (‘MeS’) derivatives of chaetocin and chetomin had no activity relative to parent compounds handled in parallel. Taken together, the data in Figure 5 indicate that dimeric chaetocin and chetomin induce *numr-1/2* and that ETP warheads are required.

### numr-1/2 is required for resistance to chetomin

Robust induction raises the possibility that *numr-1/2* may be an adaptive response to dimeric ETPs. Using an LP Sampler and BIOSORT flow-cytometer, we quantified relative length of worms added to a range of chaetocin and chetomin concentrations at the initial L1 larval stage and grown for two days. As shown in Figure 6A, larval growth is resistant to chaetocin up to at least 12.5 μM and loss of *numr-1/2 via* a CRISPR Cas9 deletion had only a minor (6%) effect on worm length at 12.5 μM. Alternatively, chetomin reduced length of wild type worms by 28% at 12.5 μM; *numr-1/2* mutants were more sensitive with length reduced by 61% in 12.5 μM chetomin. As shown in Figure 6C, an extrachromosomal array carrying *numr-1* and *numr-2* gDNA fragments fully rescued length of *numr-1/2* mutants in chetomin to the same levels as wild type worms. We obtained similar results from a second pair of independent extrachromosomal array lines derived from the same injection mix (Fig. S5C). Extrachromosomal arrays carrying *numr-1* and *numr-2* gDNA from the same injection mix had no effect on chetomin sensitivity in wild type worms (Figs. 6C and S5C). Gliotoxin, which has minimal effect on *numr-1p::GFP* (Fig. 5), strongly reduced growth of wild type and *numr-1/2* worms by similar levels at 50 μM (Fig. 6D). Taken together, these data indicate that *numr-1/2* is required for resistance to dimeric ETP chetomin, but not ETP monomer gliotoxin.

**Figure 6.**
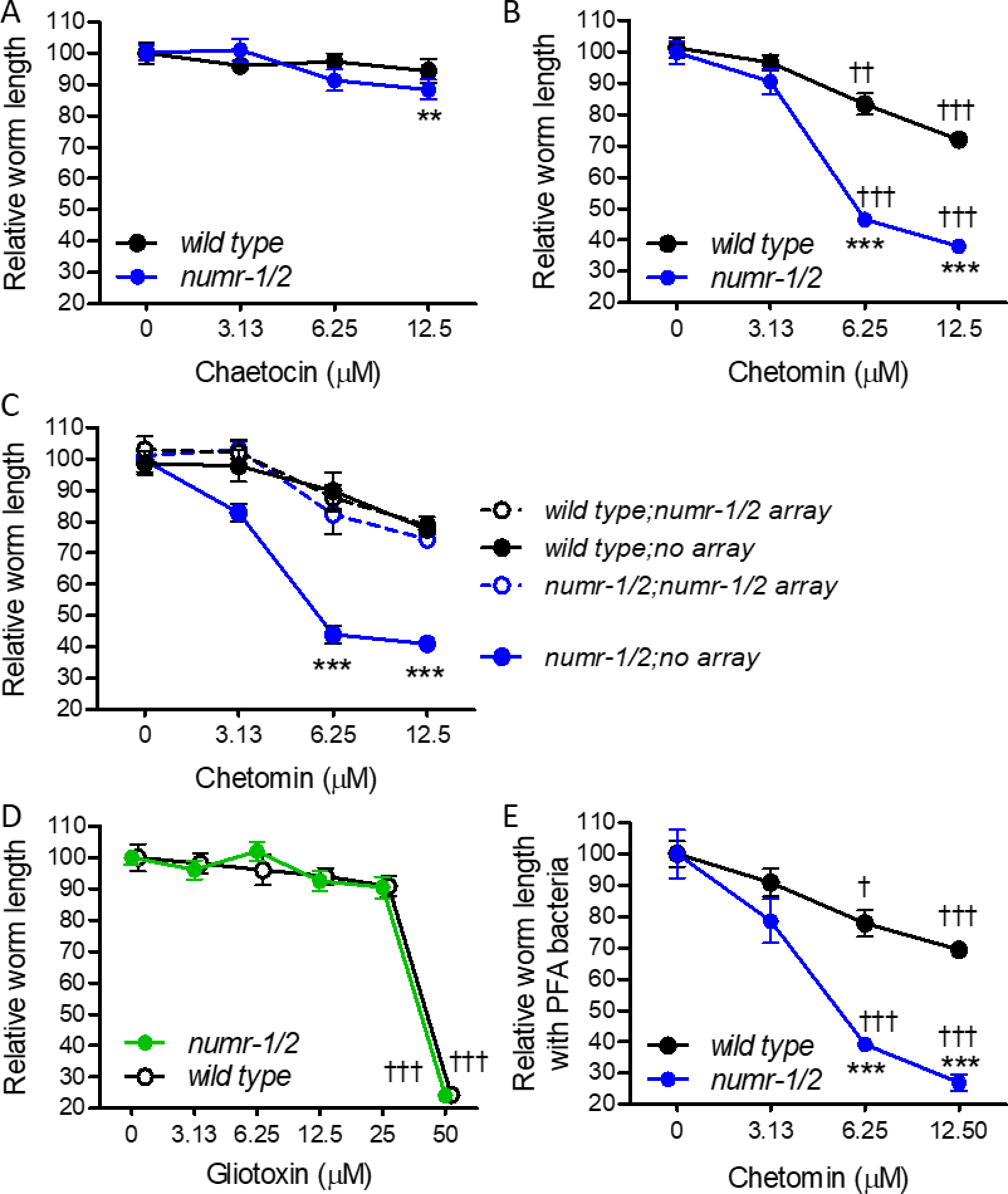
*numr-1/2* mutants are sensitive to chetomin. (A-B) Relative length of wild type and *numr-1/2* deletion worms after two days of growth with chaetocin or chetomin. N = 6 replicate wells of 29-106 worms per well from three trials for each dose. Mean ± standard error. \*\**p* < 0.01 and \*\*\**p* < 0.001 versus wild type worms at the same dose. ††*p* < 0.01 and †††*p* < 0.01 versus vehicle controls. (C) Relative length of wild type and *numr-1/2* deletion worms carrying extrachromosomal arrays with *numr-1* and *numr-2* gDNA fragments after two days of growth with chetomin. Fluorescence of co-injection marker *myo-2p::tdTomato* was used to distinguish worms with and without arrays in each well. N = 6 replicate wells of 17-50 worms per well from two trials for each dose. Mean ± standard error. \*\*\**p* < 0.001 versus wild type worms without a *numr-1/2* gDNA array at the same dose. (D) Relative length of wild type and *numr-1/2* deletion worms after two days of growth with gliotoxin. N = 6 replicate wells of 23-138 worms per well from three trials for each dose. (E) Relative length of wild type and *numr-1/2* deletion worms after two and a half days of growth with paraformaldehyde (PFA) treated bacteria and chetomin. N = 5 replicate wells of 30-104 worms per well from or one trial for each dose. (D-E) Mean ± standard error. \*\*\**p* < 0.001 versus wild type worms at the same dose. †*p* < 0.05 and †††*p* < 0.01 versus vehicle controls.

Unlike our reporter assays that started after larval development was complete, growth assays require live bacteria because early larvae either do not develop or grow very slow when raised on heat-killed bacteria or axenic media^72^. Recent studies demonstrated that bacteria treated with a low dose of paraformaldehyde do not divide and have no metabolism as measured by oxygen consumption but support worm development and size nearly as well as live bacteria^30–31, 73^. We tested this new diet model on worms grown in chetomin; as shown in Figure 6E, results for wild type and *numr-1/2* deletion worms were very similar to live bacteria in Figure 6B. These results indicate that the effects of chetomin on growth and the role of *numr-1/2* are not dependent on live bacteria.

### Chaetocin and chetomin can suppress mtl-2 promoter activity independent of numr-1/2

As shown above, 12.5 μM chaetocin suppresses *mtl-2* metallothionein gene promoter activity and mRNA levels. We next tested the effects of 3.125 μM chetomin, a dose that causes robust *numr-1p::GFP* induction (Fig. 5C) on *mtl-2p::GFP* and found a 44% suppression similar to 12.5 μM chaetocin in the same trials (41%) (Fig. 7). We next crossed the *mtl-2p::GFP* reporter with the *numr-1/2* deletion allele used in Figure 6. As shown in Fig. 7, 12.5 μM chaetocin and 3.125 μM chetomin suppressed *mtl-2p::GFP* in the *numr-1/2* deletion mutant by 30 and 39%, respectively; these effects were statistically reduced relative to wild type, but still robust. Therefore, *numr-1/2* deletion may influence regulation of the *mtl-2* promoter, but most of the repression by chaetocin and chetomin can occur independently.

**Figure 7.**
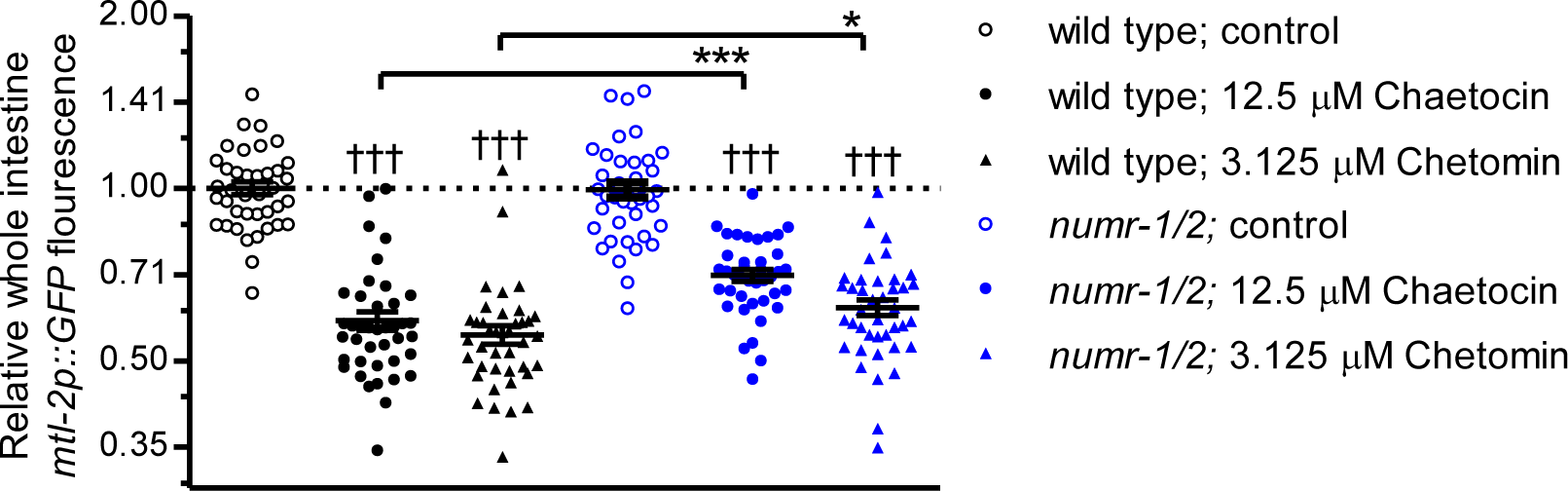
Chaetocin and chetomin are able to suppress *mtl-2p::GFP* in *numr-1/2* mutants. Scatter plot of relative *mtl-2p::GFP* fluorescence with chaetocin or chetomin in wild type or *numr-1/2* deletion backgrounds. Chaetocin and chetomin suppressed *mtl-2p::GFP* fluorescence in both strains. Mean ± standard error. †††*p* < 0.01 versus vehicle controls; \**p* < 0.05 and \*\*\**p* < 0.001 for *numr-1/2* versus wild type worms. N = 39-40 individual worms from two trials for each condition.

### Chaetocin and chetomin affect expression of stress-response genes in human cells

Lastly, we performed RNAseq to explore the possibility of chaetocin and chetomin affecting expression of genes related to metal homeostasis in a cell line derived from human intestine. HCT116 colorectal cancer cells, used as model system, were exposed to 500 nM chaetocin, a dose that blocks proliferation^59, 74^, and RNA was isolated after 5 and 12 h to identify early and late expression changes. Expression was also measured for the same dose of chetomin and methylated sulfide (‘MeS’) analogs. A heat map of all relative expression changes for all conditions is shown in Figure 8A. Chaetocin and chetomin had correlation coefficients of 0.890 at 5 h and 0.916 at 12 h indicating very similar effects, and expression changes were almost completely dependent on disulfides in the ETP cores (Fig. 8A).

**Figure 8.**
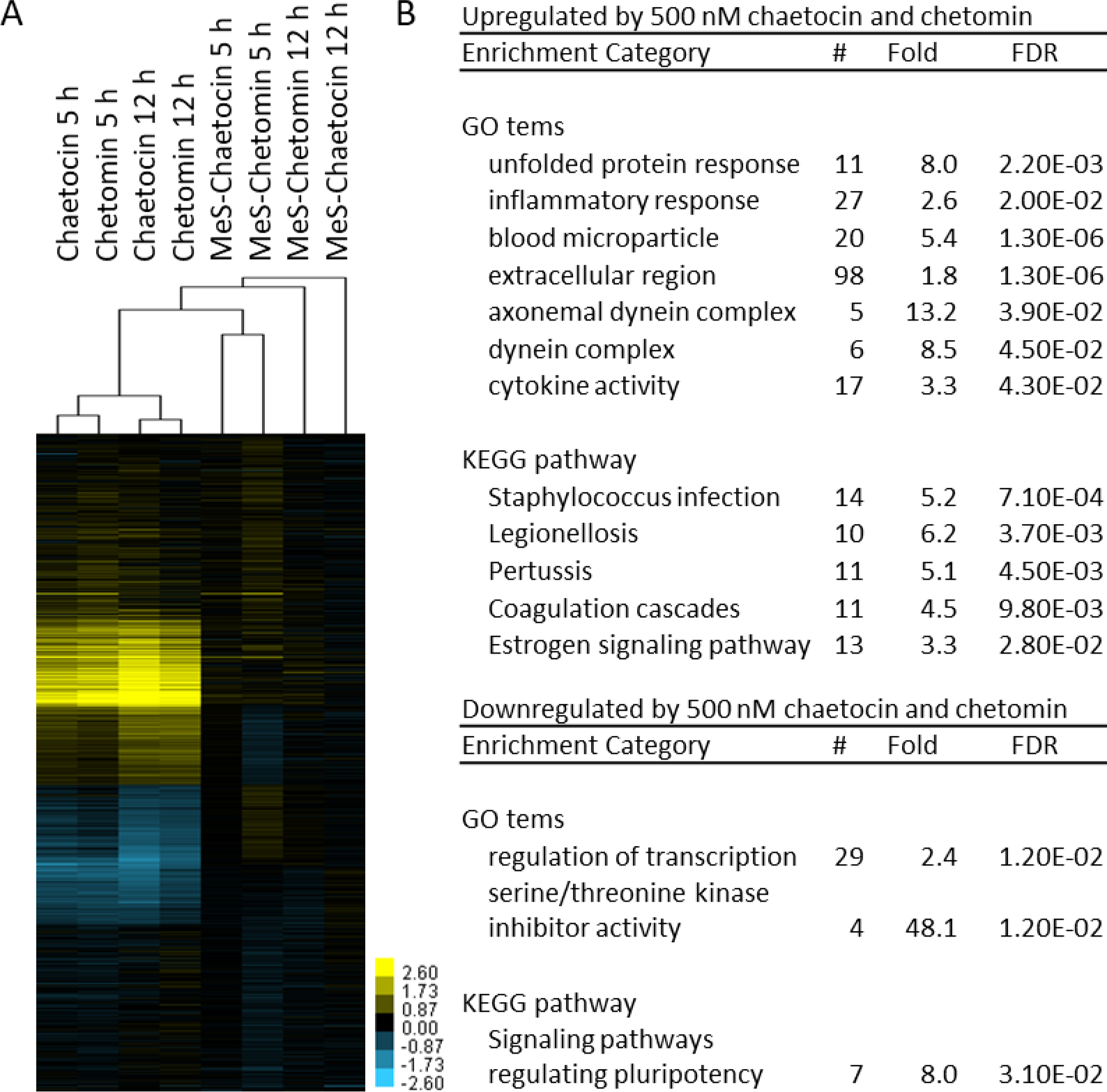
Chaetocin and chetomin affect expression of human genes related to immune response and hemostasis. (A) Heat map of mean relative expression changes for all conditions clustered by gene and condition (log2). (B) Gene ontology (GO) and KEGG terms enriched among genes affected at least 2-fold by chaetocin and chetomin at both time points but by not methylated analogs; FDR, false discovery rate.

To generate high-confidence lists, we identified genes affected at least 2-fold by chaetocin and chetomin at both time points and subtracted any genes also affected by the MeS analogs; this yielded 818 genes upregulated and 160 genes downregulated by both intact ETPs at both time points (Table S6). The top categories for upregulated genes were related to unfolded protein response, inflammation and infection responses, and hemostasis (Fig. 8B). We also looked closely at HCT116 data for genes with well-characterized roles in cellular stress responses. Chaetocin and chetomin, but not MeS analogs, increased expression of several redox, heat shock, protein misfolding, and osmotic stress response genes consistent with broad activity in cells including redox imbalance (Fig. S7)^57, 75^. Interestingly, three metallothionein genes (MT1E, MT1X, and MT2A) and a metal ion transport gene (SLC39A7) were downregulated by chaetocin and chetomin at 12 h (Fig. S6).

## DISCUSSION

Using a chemical genetic approach, our study links fungal ETPs chaetocin and chetomin to a novel nuclear metal response. Chaetocin and chetomin are distinct from cadmium for their ability to induce *numr-1/2* and repress metallothionine genes. Our results in Fig. 7 indicate that the mechanism of *mtl-2* promoter suppression does not require *numr-1/2* as most activity is retained in a deletion mutant.

In mammals, zinc-finger protein MTF-1 (metal response element-binding transcription factor 1) activates metallothionein gene expression during exposure to cadmium by recruiting transcriptional co-factors to promoters^76^. In *C. elegans,* cadmium was recently shown to directly bind to nuclear receptor HIZR-1 (high zinc activated nuclear receptor) as a ligand to activate expression of metallothionein genes^42, 77^. *C. elegans* has additional metal-responsive signaling mechanisms because *numr-1/2* and many other genes can be strongly activated by cadmium independently of HIZR-1^42^. Repression of metallothionein genes by chaetocin and chetomin is also consistent with a distinct mechanism (Figs. 3, 7, S3, and S5)^77^.

Redox imbalance has been measured in cancer cells exposed to ETPs^16, 19, 78^, which is consistent with activation of HMOX1 and unfolded protein responses that we observed in HCT116 cells (Fig. S6). In *C. elegans*, canonical redox and unfolded protein response genes (e.g., *gst-4* and *hsp-4*) were not activated by chaetocin or chetomin (Figs. S3 and S5) and loss of thioredoxin reductases had no effect on *numr-1/2* expression (Fig. S4). Chaetocin and chetomin block binding of the CH1 domain of p300 to transcription factor HIF-1α *in vitro* and in human cells^59, 79^. RNAi of the single *C. elegans* p300/CBP orthologue *cbp-1* did not induce *numr-1p::GFP* in the alimentary canal like chaetocin or chetomin (Fig. S4D) implicating a different mechanism. Our prior genome-wide screen and current RNAi screen did not identify any predicted histone methyltransferases with effects on *numr-1p::GFP*. However, histone methyltransferases are a large protein family in *C. elegans,* and further work is needed to determine if they play a role in chaetocin or chetomin responses^22–23, 71^. Future unbiased screens for genes required for *numr-1/2* activation could identify new chaetocin or chetomin targets and mechanisms of metal-response gene regulation.

Cook et al., previously demonstrated that an ETP disulfide core is required for activity against p300 *in vitro*^59^. We found that ETP disulfides were also required for chaetocin and chetomin to induce *numr-1p::GFP* in *C. elegans* (Fig. 5) and to induce gene responses in HCT116 cells (Fig. 8). Cook et al., also demonstrated that monomer gliotoxin had similar activity as chaetocin and chetomin suggesting that an ETP disulfide core was sufficient^59^. In contrast, gliotoxin has little to no activity on *in vivo numr-1p::GFP* despite having other robust bioactivities^68–69^ (Figs. 5-6). Accessibility to organelles and target proteins is likely to vary depending on the structure of compounds containing ETPs. Understanding the structural features of chaetocin and chetomin that distinguish their ability to activate *numr-1/2* compared to gliotoxin could provide insights into how this versatile class of molecules can be tailored to specific targets.

Interest in ETPs is growing, and variants designed for derivatization have been synthesized that retain activity against a wide range of cancer cell types^57, 80–81^. ETP producing fungi infect immunocompromised individuals and mycotoxins are environmental hazards present in damp buildings and food crops^82–83^. While mechanisms for ETP toxicity have been identified, very little is known about adaptive responses that promote resistance in cells other than in the Chaetomiaceae fungi that synthesize them^84^. Understanding these responses could help improve safety and efficacy, provide insights into how cells might develop resistance, and improve treatment of infections and exposures. Chaetomiaceae fungi and *C. elegans* inhabit similar environments raising the possibility that *numr-1/2* may function to help nematodes cope with ETP mycotoxins in nature. Biochemical functions of NUMR-1/2 are unclear, but the protein is expressed in nuclei of alimentary canal cells that contact the environment and contains an RNA recognition motif and histidine-rich domain^4, 6^. The RNA recognition motif of NUMR-1/2 could facilitate interactions with nucleic acids. Histidine-rich domains in other proteins reversibly bind zinc, small molecules, and other proteins raising the possibility of a direct role in metal homeostasis, signaling, or interactions with ETPs^2, 11, 85–86^. There is no direct homolog of NUMR-1/2 in human cells, but chaetocin and chetomin did induce gene HRG (histidine-rich glycoprotein) 32-fold (Fig. S6); HRG protein has many functions including zinc chaperone and small molecule binding^2, 85–86^. Future studies testing binding to metals and ETPs could help define NUMR-1/2 biochemical functions and mechanisms of resistance.

Modulation of zinc signaling could explain broader effects of chaetocin and chetomin on gene expression that we observed in HCT116 cells. Zinc binds to about 10% of the human proteome acting as a co-factor for enzymes and transcriptional regulators^87^. Genes activated by chaetocin and chetomin in HCT116 cells are enriched for functions in infection and hemostasis (Fig. 8), and similar results were recently reported for esophageal squamous cell carcinoma cells exposed to chaetocin^88^. Several core immune response signaling proteins bind zinc, which is considered a “gate-keeper” for immune function^87^. Zinc also has numerous roles in hemostasis^89^. Displacement of zinc-binding from proteins was proposed as a mechanism for chetomin in HCT116 cells^59^. Zinc may play a different role in activation of *numr-1/2* in *C. elegans* because zinc chelator TPEN had little effect on its own and did not repress chaetocin activity (Fig. 4). These results suggest that the relationships between ETPs and zinc may be nuanced and target-specific.

## ACKNOWLEDGEMENTS

We would like to acknowledge P. Kamble and C. Huss for assistance with compound library dispensing for the screen. Some strains were provided by the *Caenorhabditis* Genetics Center, which is funded by NIH Office of Research Infrastructure Programs (P40 OD010440). The BIOSORTER used for screening was obtained with NIH grant S10-OD012006.

