## Supporting information for "Chemical genetics in *C. elegans* identifies anticancer mycotoxins chaetocin and chetomin as potent inducers of a nuclear metal homeostasis response"

Supplementary gene expression, gene reporter, and growth data and experimental details for compound modification are provided as supporting information as follows:

Figures:

Figure S1. Structures of *numr-1p::GFP* inducers and chaetocin analogs tested

Figure S2. Inducers activate *numr-1p::GFP* in the alimentary canal or late embryos

Figure S3. RT-qPCR of *numr-1/2* and representative stress response genes with chaetocin and tannic acid

Figure S4. Effects of *trxr-1/2* or *cbp-1* loss on *numr-1/2*

Figure S5. Chetomin activates *numr-1/2* and *numr-1/2* promotes resistance

Figure S6. Chaetocin and chetomin affect stress-response genes in HCT116 cells

Preparation methods for MeS-chaetocin and MeS-chetomin

Tables:

Table S1. List of screened libraries, number of compounds tested per library, and test concentrations

Table S2. List of primers

Table S3. Details of candidate compounds that activated *numr-1p::GFP*

Table S4. List of methyltransferase dsRNA clones in siRNA library tested

Table S5. $^{1}H$ NMR data for S-methyl-products of chaetocin and chetomin

Table S6. List of humans genes up- and down-regulated by 500 nM chaetocin and chetomin

### Figure S1

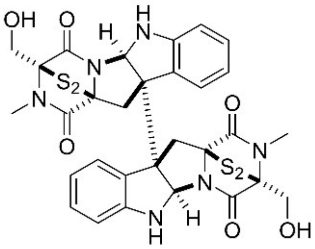

Chaetocin

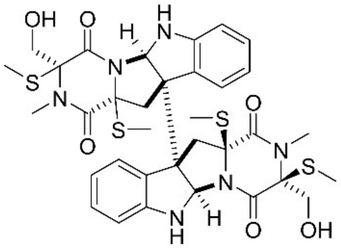

MeS-Chaetocin

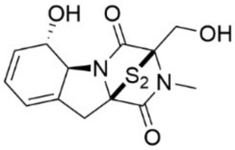

Gliotoxin

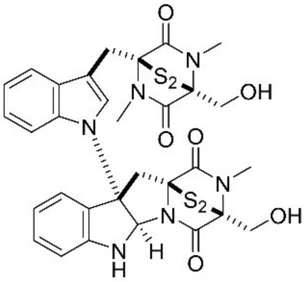

Chetomin

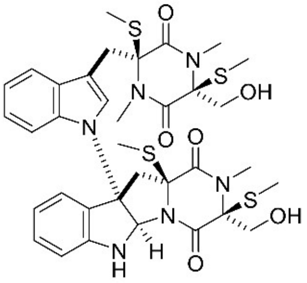

MeS-Chetomin

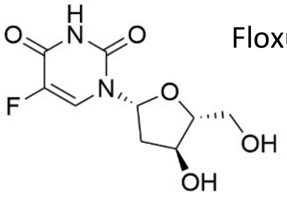

Floxuridine

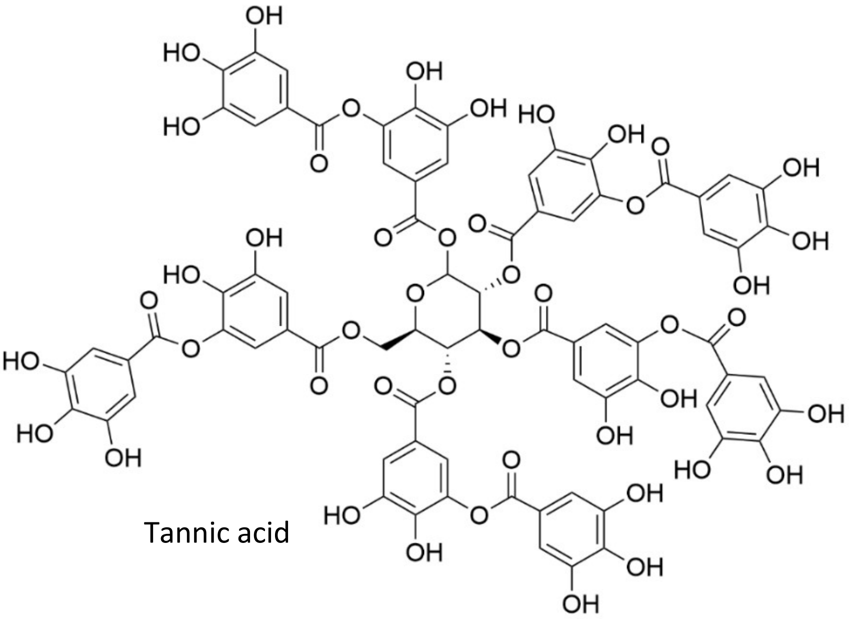

Tannic acid

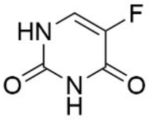

5-Fluorouracil

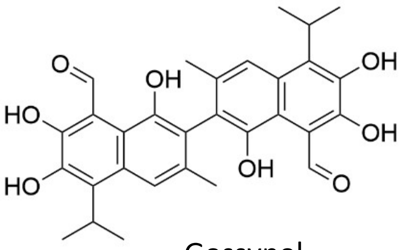

Gossypol

Figure S1. Structures of *numr-1p::GFP* inducers and chaetocin analogs tested.

### Figure S2

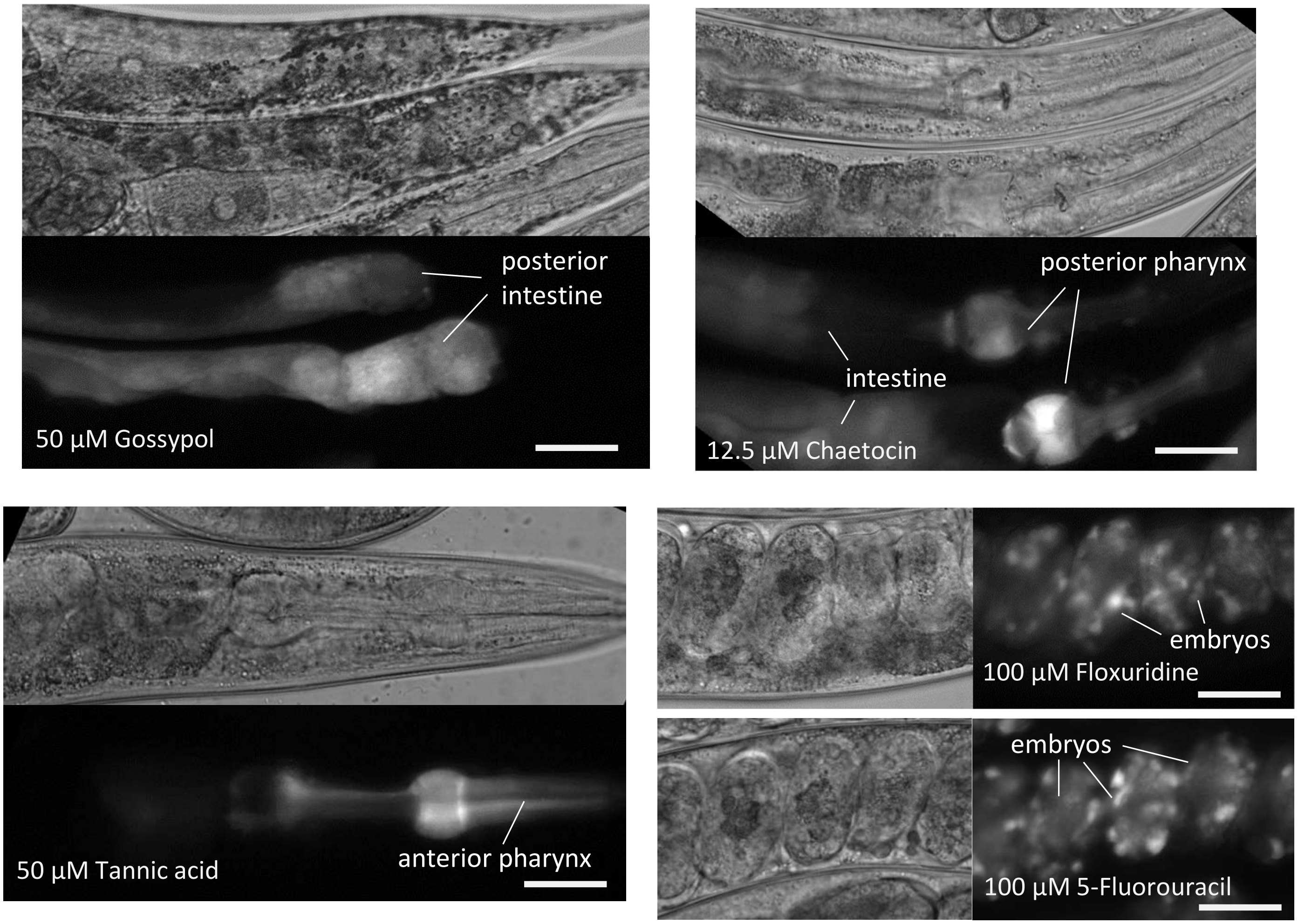


Figure S2. Inducers activate *numr-1p::GFP* in the alimentary canal or late embryos. Bright field and *numr-1p::GFP* micrographs are paired. All scale bars are 25 µm.

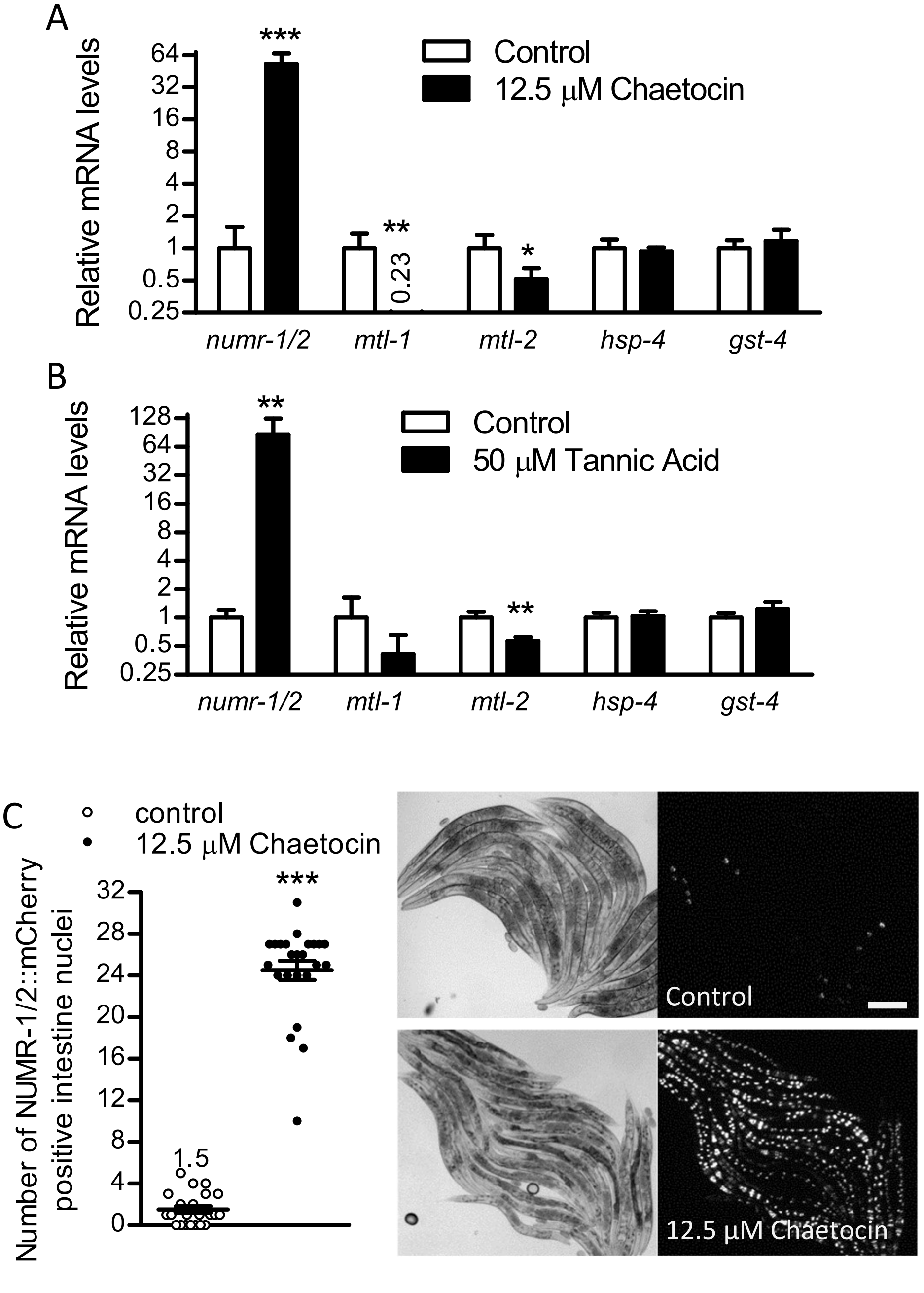


Figure S3. RT-qPCR confirms activation of *numr-1/2* by chaetocin and tannic acid. (A-B) Expression of *numr-1/2* and representative stress response genes was measured with RT-qPCR in worms exposed to chaetocin or tannic acid for 5 h. (A-B) Mean + standard error, N = 5 cDNA samples of 10 worms each. *$p < 0.05$, **$p < 0.01$, and ***$p < 0.001$. (C) Number of intestinal nuclei with visible NUMR-1::mCherry fluorescence per worm (left) and bright field NUMR-1::mCherry micrographs (right); scale bar is 200 μm. Mean ± standard error. N = 22-24 individual worms per condition from one trial. (A-C) *$p < 0.05$, **$p < 0.01$, and ***$p < 0.001$ for treatments versus vehicle controls.

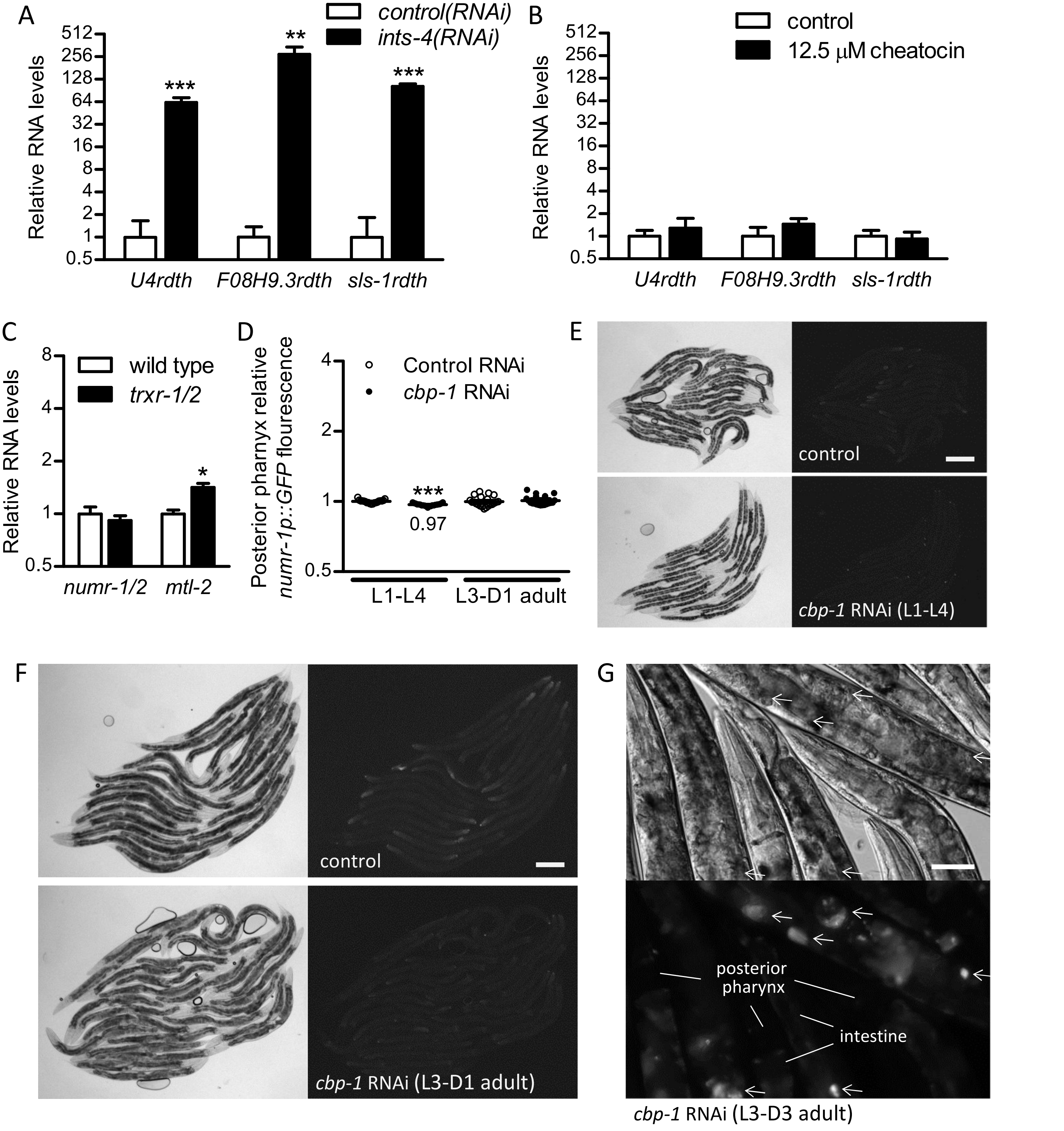


Figure S4. Effects of *trxr-1/2* or *cbp-1* loss on *numr-1/2*. (A-C) Relative RNA levels measured by RT-qPCR. (A-B) RNAi of integrator subunit *ints-4* strongly increases levels of read-through (rdth) transcripts, but 12.5 μM chaetocin does not. Mean + standard error. (A) N = 5-8 cDNA samples of 5 worms each. (B) N = 5 cDNA samples of 10 worms each. (C) Deletion of *trxr-1* and *trxr-2* does not mimic the effects of chaetocin on expression of *numr-1/2* and *mtl-2*. Mean + standard error, N = 5 cDNA samples of 10 worms each. (D) Relative *numr-1p::GFP* fluorescence. Mean ± standard error normalized to control RNAi within stage. N = 19-21 individual worms per condition from one trial. (E-G) Bright field and *numr-1p::GFP* micrographs in worms treated with RNAi from L1 to L4 larval stage (E), L3 to day 1 adulthood (F), or L3 to day 3 adulthood (G, signal marked with arrowheads); scale bars are 200 μm (E-F) or 50 μm (G). (A-D) *$p < 0.05$, **$p < 0.01$, and ***$p < 0.001$ for treatments versus vehicle, wild type, or RNAi controls.

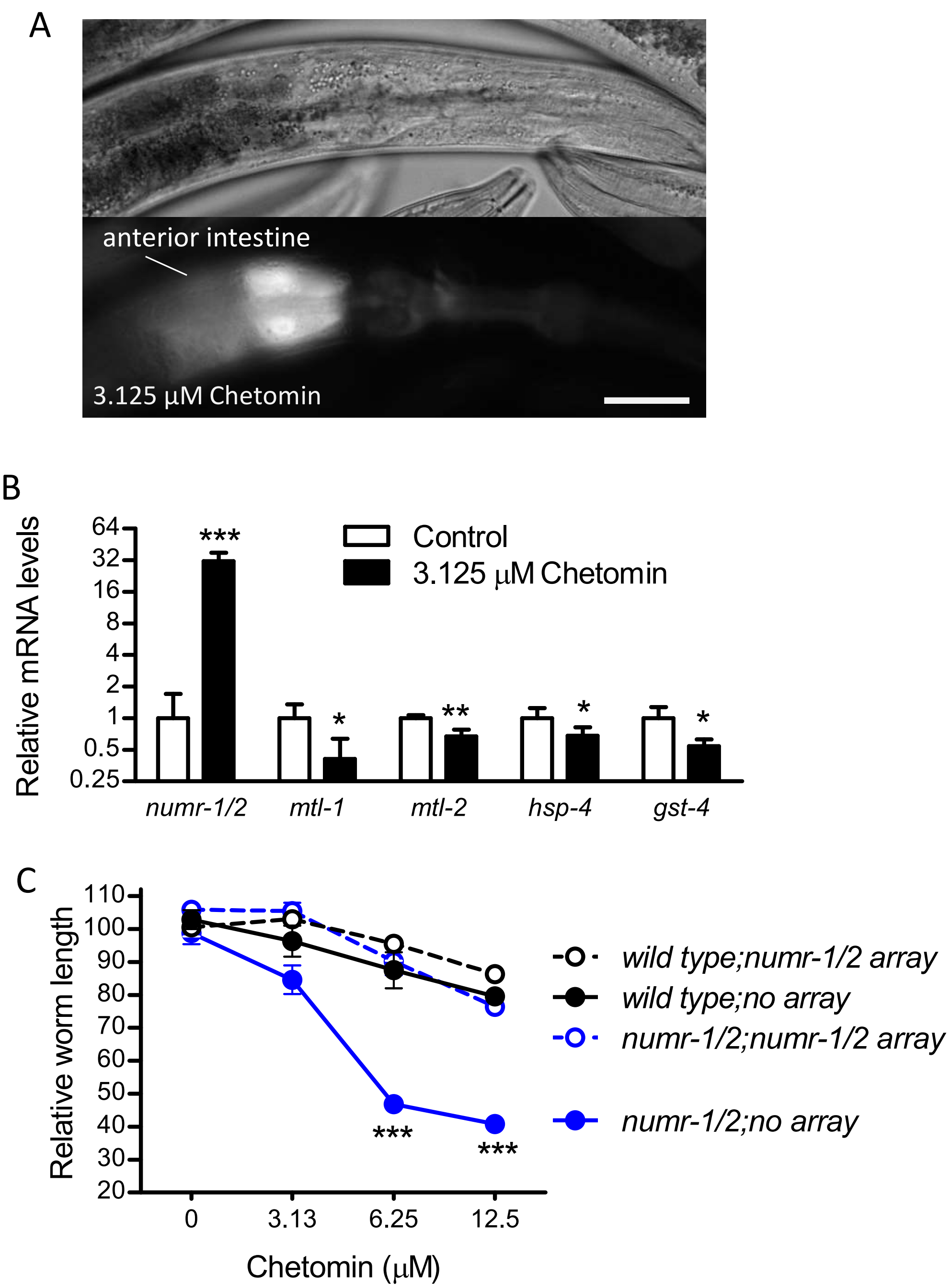


Figure S5. Chetomin activates *numr-1/2* and *numr-1/2* promotes resistance. (A) Bright field and *numr-1p::GFP* fluorescence micrographs. Scale bar is 25 μm. (B) Relative RNA levels measured by RT-qPCR. N = 4-5 cDNA samples of 10 worms each. $*p < 0.05$, $**p < 0.01$, and $***p < 0.001$ for treatment versus vehicle controls. (C) Relative length of wild type and *numr-1/2* deletion worms carrying extrachromosomal arrays with *numr-1* and *numr-2* gDNA fragments after two days of growth with chetomin. Fluorescence of co-injection marker *myo-2p::tdTomato* was used to distinguish worms with and without arrays in each well. N = 5 replicate wells of 16-101 worms per well from two trials for each dose. Mean ± standard error. $***p < 0.001$ versus wild type worms without a *numr-1/2* gDNA array at the same dose.

Figure S6

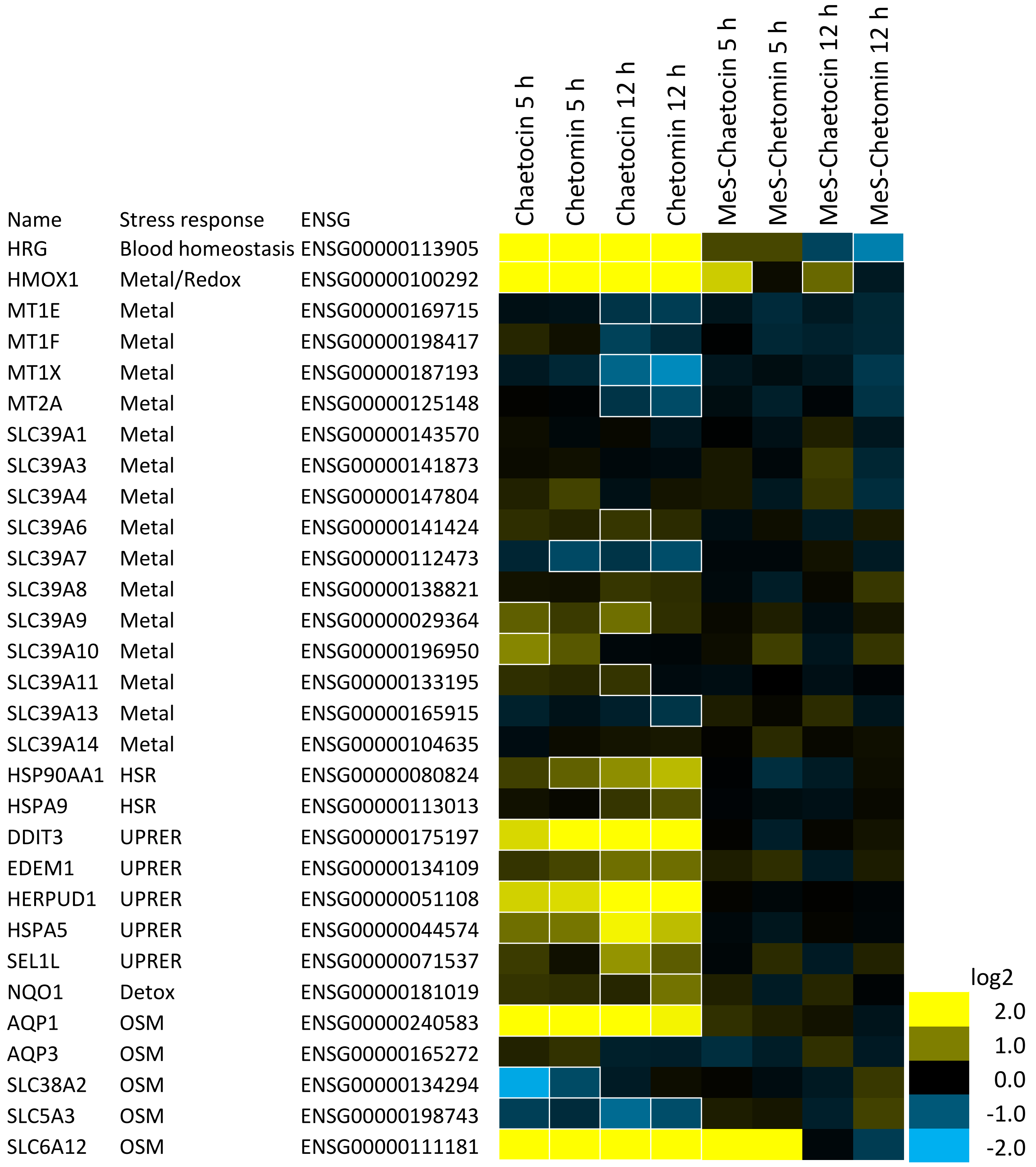


Figure S6. Chaetocin and chetomin affect stress-response genes in HCT116 cells. Heat map of mean relative expression changes at 5 h and 12 h from HCT116 RNAseq data. Significant changes are outlined in a white boarder. Many protein misfolding response genes were upregulated and metal-response genes were downregulated. HSR (heat shock response), UPRER (unfolded protein response), Detox, (detoxification response), OSM (osmotic stress response).

*Preparation of MeS-chaetocin and MeS-chetomin*

MeS-chaetocin and MeS-chetomin are structural analogues of commercially available chaetocin and chetomin, in which the sulfur bridges have been broken and alkylated. Commercial parent compounds were purchased from Cayman Chemical. The scheme for modification, structure data, and supplementary references are shown below.

*Reduction of chaetocin and chetomin to S-methyl-products*

Pyridine (75 µL), MeI (50 µL) and $NaBH_4$ (2.2 mg, 0.058 mmol) were successively added to a solution of chaetocin or chetomin (5 mg, 0.007 mmol) in MeOH (125 µL). The reaction mixture was stirred at room temperature for 5 h under an argon atmosphere. The reaction was quenched by brine, extracted with ethyl acetate, dried with anhydrous $MgSO_4$, and concentrated *in vacuo* to give crude product, which was further purified by HPLC to afford pure S-methyl-products, MeS-chaetocin, and MeS-chetomin. Briefly, crude mixtures of containing MeS-chaetocin and MeS-chetomin were purified via reversed-phase HPLC (Phenomenex Synergi Fusion-RP; 250 X 10mm; 4µm) using a linear $H_2O$-MeCN gradient (0-100% MeCN in 10 min, 100% MeCN for 10 min). MeS-chaetocin ($t_R$, 14.6 min) and MeS-chetomin ($t_R$, 15.9 min) structures were confirmed by 600 MHz $^1$H NMR and mass spectrometry and compared to data reported in the literature (Table S6).

*NMR conditions used for structure determination of MeS-chaetocin and MeS-chetomin*

MeS-chaetocin: 1.4 mg, $[\alpha]_D$: +92.5 (*c* 0.0004, $CHCl_3$, 21 °C). $^1$H NMR in Table S5 (600 MHz, $CDCl_3$): HRESIMS m/z 779.1771 $[M+Na]^+$ (calcd for $C_{34}H_{40}O_6N_6NaS_4$ 779.1784).

MeS-chetomin or Dethio-tetra (methylthio) chetomin: 1.2 mg, $[\alpha]_D$: +66.0 (*c* 0.0004, $CHCl_3$, 21 °C). $^1$H NMR in Table S6 (600 MHz, $CDCl_3$): HRESIMS m/z 793.1941 $[M+Na]^+$ (calcd for $C_{35}H_{42}O_6N_6NaS_4$ 793.1941).

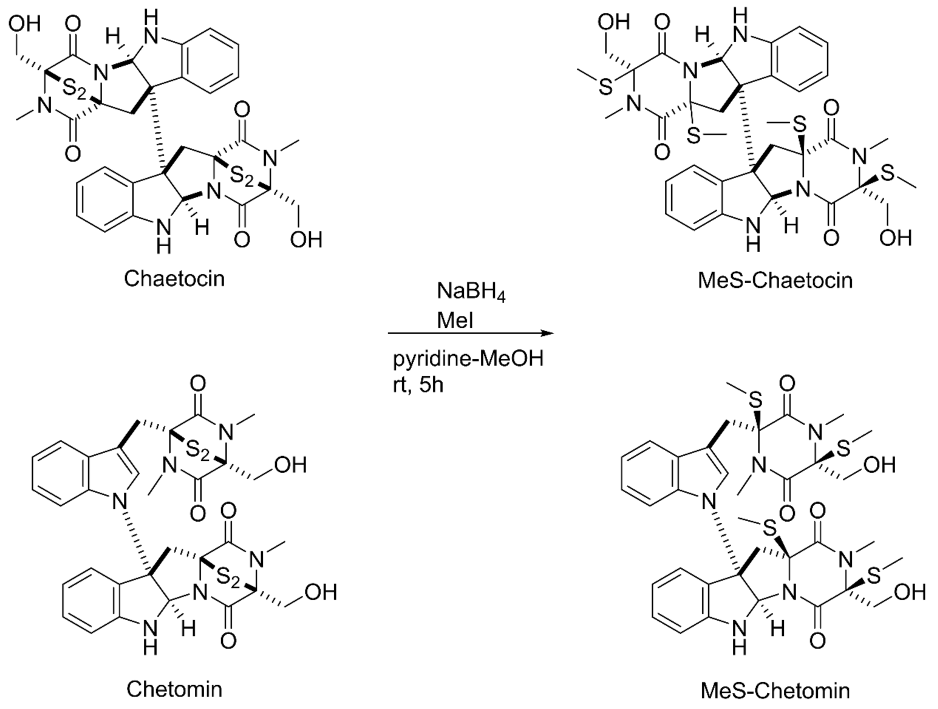

Table S1 Libraries

| Library | Source | Compounds | Concentrations |
|---|---|---|---|
| Epigenetic Small Molecule Library | Cayman Chemicals | 145 | 50, 10. 1. 0.1 µM |
| LOPAC | Sigma, Millipore | 1280 | 50 µM |
| Spectrum | Microsource Discovery Systems, Inc. | 2000 | 50 µM |
| Prestwick | Prestwick | 1520 | 50 µM |
| NCI Program for Natural Product Discovery | NIH NCI | 36771 | 10 µg/ml |

Table S2 Primers

| qPCR primer | Sequence |
|---|---|
| rpl-2 F | CTTTCCGCGACCCATACAA |
| rpl-2 R | CACGATGTTTCCGATTTGGAT |
| gst-4 F | TCCGTCAATTCACTTCTTCCG |
| gst-4 R | AAGAAATCATCACGGGCTGG |
| hsp-4 qPCR F' | GGATCAACCAGAATTCCAAAGG |
| hsp-4 qPCR R' | TCAGGGTTGATTCCACGAGAT |
| mtl-2 F3 | ATGGTCTGCAAGTGTGACTGC |
| mtl-2 R3 | GGCAGTTGGGCAGCAGTATT |
| mtl-1 F | CATGGCTTGCAAGTGTGACTG |
| mtl-1 R | TCTCACTGGCCTCCTCACAG |
| numr-1 qF1 | AGACGTCACTGTTTTGGTGGA |
| numr-1 qR1 | CCGAATCCTCCAGTTGGACC |
| U4 Misproesssing F | CCCCTGAAACATGGGTGGCATACG |
| U4 Misproesssing R | ACGCGATTACGGTGTAATGCCAGA |
| F08H9.3 F | CGCATGGGCTCTCTGGTAAA |
| F08H9.3 R | TCGAAAAACGGGTTGACGGA |
| sls-1 F | GATCCCGTCCCCCAATCAAT |
| sls-1 R | CCATGTTGGAGAACGGTCCA |

| gDNA primer | Sequence |
|---|---|
| numr-1 F1 | CGCTCTCCAGATTTCAGACCC |
| numr-1 R1 | TGAAGATTCGGCTGCCAGAG |
| numr-2 F1 | TTCCAAACATCCCATCCCGA |
| numr-2 R2 | GGACTGAGTGCCTGCCTATG |

Table S3 Compounds

| Name | Library | EC50 | 50 µM | 10 µM | 1 µM | 0.1 µM |
|---|---|---|---|---|---|---|
| Floxuridine | Spectrum | 32.5 µM | + | ND | ND | ND |
| Floxuridine | Prestwick | 32.5 µM | + | ND | ND | ND |
| Gossypol | Spectrum | 26.0 µM | + | ND | ND | ND |
| Tannic acid | Spectrum | 17.1 µM | + | ND | ND | ND |
| Chaetocin | Epigenetic Small Molecule Library | 5.1 µM | + | + | + | - |
| Pyrromycin | Spectrum | Not tested | + | ND | ND | ND |
| 2',2'-bisepigallocatechin digallate | Spectrum | Not tested | + | ND | ND | ND |
| SP600125 | LOPAC | not confirmed | + | ND | ND | ND |
| JIB-04 | Epigenetic Small Molecule Library | not confirmed | + | - | - | - |
| Berberine chloride | Spectrum | not confirmed | + | ND | ND | ND |
| Actinomycin D | Spectrum | toxic | + | ND | ND | ND |
| 5-Fluorouracil | Spectrum | 53.7 µM | + | ND | ND | ND |
| 5-Fluorouracil | Prestwick | 53.7 µM | + | ND | ND | ND |
| 4-Chloromercuribenzoic acid | LOPAC | Not tested | + | ND | ND | ND |
| Toltrazuril | Spectrum | Not tested | + | ND | ND | ND |
| 2,3-Dimercaptosuccinic acid | Spectrum | Not tested | + | ND | ND | ND |
| 2,3-Dichloro-5,8-dihydroxynapthoquinone | Spectrum | Not tested | + | ND | ND | ND |
| 3alpha-acetoxydihydrodeoxygedunin | Spectrum | Not tested | + | ND | ND | ND |
| Theaflavin monogallates | Spectrum | Not tested | + | ND | ND | ND |
| Epicatechin monogallate | Spectrum | Not tested | + | ND | ND | ND |

+ (activated numr-1/2p::GFP), - (no activation), ND (not determined)

Table S4 Histone methyltransferases

| | |
|---|---|
| mes-4 | Y2H9A.1 |
| met-2 | R05D3.11 |
| set-1 | T26A5.7 |
| set-10 | F33H2.7 |
| set-11 | F34D6.4 |
| set-12 | K09F5.5 |
| set-14 | R06F6.4 |
| set-15 | R11E3.4 |
| set-16 | T12D8.1 |
| set-17 | T21B10.5 |
| set-18 | T22A3.4 |
| set-2 | C26E6.9 |
| set-20 | W01C8.4 |
| set-22 | Y32F6A.1 |
| set-23 | Y41D4B.12 |
| set-24 | Y43F11A.5 |
| set-25 | Y43F4B.3 |
| set-26 | Y51H4A.12 |
| set-3 | C07A9.7 |
| set-30 | ZC8.3 |
| set-32 | C41G7.4 |
| set-4 | C32D5.5 |
| set-5 | C47E8.8 |
| set-9 | F15E6.1 |

**Table S5**. NMR (600 MHz in $CDCl_3$) data for S-methyl- Chaetocin and Chetomin

| | Chaetocin† | MeS-chaetocin | | Chetomin‡ | MeS-chetomin | |
|---|---|---|---|---|---|---|
| C/H no. | $\delta_H$ (*J* in Hz) | $\delta_H$ (*J* in Hz) | $\delta_C$ | $\delta_H$ (*J* in Hz) | $\delta_H$ (*J* in Hz) | $\delta_C$ |
| 1 | - | - | - | - | - | 167.6 |
| 2 | 5.16, s | 5.74, s | 81.3 | 3.20, s | 3.22, s | 32.6 |
| 3 | - | - | - | - | - | 73.9 |
| 3-$CH_2$ | - | - | - | 4.31, d (12.7)<br>4.38, d (12.7) | 3.86, m<br>4.39, d (11.7) | 66.2 |
| 3-OH | - | - | - | - | - | - |
| 3-S$CH_3$ | - | - | - | - | 2.27, s | 13.8 |
| 4 | - | - | 129.6 | - | - | 168.5 |
| 5 | 6.64, d (7.8) | 6.64, d (7.8) | 110.9 | 6.21, s | 6.17, s | 83.7 |
| 6 | 6.82, dd (7.6) | 6.87, dd (7.4) | 119.9 | - | - | 131.9 |
| 7 | 7.16, dd (7.5) | 7.21, dd (7.6) | 130.6 | 6.81, d (7.9) | 6.74, d (7.9) | 112.3 |
| 8 | 7.35, d (7.5) | 7.49, d (7.4) | 126.1 | 7.31, m | 7.27, m | 133.0 |
| 9 | - | - | 151.6 | 6.96, t (7.5) | 6.86, t (7.4) | 121.9 |
| NH | - | 4.87, d (3.2) | - | 5.30, s | 5.11, brs | - |
| 10 | - | - | - | 7.35, d (7.2) | 7.29, m | 126.6 |
| 10a | - | - | - | - | - | 133.9 |
| 10b | - | - | - | - | - | 71.2 |
| 11 | - | - | 69.7 | 3.11, d (15.4)<br>4.43, d (15.5) | 3.38, d (14.3)<br>3.60, d (14.3) | 46.8 |
| 11a | - | - | - | - | - | 84.4 |
| 11-S$CH_3$ | - | 1.87, s | 15.8 | - | 2.01, s | 16.5 |
| 12 | 2.69, d (15.0)<br>3.75, d (15.0) | 2.46, d (14.8)<br>2.66, d (14.8) | 44.6 | - | - | - |
| 13 | - | - | 167.1 | - | - | - |
| 14 | - | - | - | - | - | - |
| 15 | - | - | 76.2 | - | - | - |
| 15-S$CH_3$ | - | 2.19, s | 12.8 | - | - | - |
| 16 | - | - | 166.1 | - | - | - |
| 17 | 4.07, d (12.6)<br>4.19, d (12.6) | 4.31, dd (6.8, 11.6)<br>3.80, dd (7.4, 11.6) | 65.4 | - | - | - |
| 17-OH | - | 4.62, brs | - | - | - | - |
| 18 | 3.01, s | 3.10, s | 29.2 | - | - | - |
| 1’ | - | - | - | - | - | 170.1 |
| 2’ | - | - | - | 2.96, s | 2.79, s | 30.6 |
| 3’ | - | - | - | 4.28, d (12.5)<br>4.35, d (12.5) | 3.05, d (11.2)<br>3.77, dd (6.7, 11.2) | 73.9 |
| 3’-$CH_2$ | - | - | - | | | 65.9 |
| 3’-OH | - | - | - | - | - | - |
| 3’-S$CH_3$ | - | - | - | - | 2.10, s | 13.9 |
| 4’ | - | - | - | - | - | 170.6 |
| 5’ | - | - | - | 3.17, s | 3.15, s | 30.3 |
| 6’ | - | - | - | - | - | 110.3 |
| 6’-S$CH_3$ | - | - | - | - | 2.30, s | 15.1 |
| 7’ | - | - | - | 3.73, d (15.3)<br>3.89, d (15.4) | 3.21, d (14.9)<br>3.91, d (14.9) | 34.8 |
| 8’ | - | - | - | - | - | 75.7 |
| 9’ | - | - | - | 7.19, s | 7.02, s | 127.1 |
| 10’ | - | - | - | - | - | 132.1 |
| 11’ | - | - | - | 7.24, m | 7.00, m | 113.9 |
| 12’ | - | - | - | 7.24, m | 7.00, m | 124.8 |
| 13’ | - | - | - | 7.31, m | 7.07, m | 122.3 |
| 14’ | - | - | - | 7.67, d (6.9) | 7.59, d (8.0) | 121.8 |
| 14a’ | - | - | - | - | - | 137.5 |

†Consistent with published literature by (IWASA *et al.* 2010)

‡Consistent with published literature by (WANG *et al.* 2018)

Iwasa, E., Y. Hamashima, S. Fujishiro, E. Higuchi, A. Ito *et al.*, 2010 Total synthesis of (+)-chaetocin and its analogues: their histone methyltransferase G9a inhibitory activity. J Am Chem Soc 132**:** 4078-4079.

Wang, M. H., Y. C. Hu, B. D. Sun, M. Yu, S. B. Niu *et al.*, 2018 Highly Photosensitive Poly-Sulfur-Bridged Chetomin Analogues from Chaetomium cochliodes. Org Lett 20**:** 1806-1809.

Table S6 Human RNAseq

| upChaetocin&Chetomin_5h&12h_notMeS | ENSG | GeneName | downChaetocin&Chetomin_5h&12h_notM | ENSG | GeneName |
|---|---|---|---|---|---|
| | ENSG00000181409 | AATK | | ENSG00000168792 | ABHD15 |
| | ENSG00000121270 | ABCC11 | | ENSG00000279716 | AC006128.1 |
| | ENSG00000258924 | AC002094.1 | | ENSG00000232445 | AC006329.1 |
| | ENSG00000241764 | AC002467.1 | | ENSG00000268262 | AC011445.1 |
| | ENSG00000275936 | AC004263.2 | | ENSG00000273466 | AC012510.1 |
| | ENSG00000263412 | AC004477.1 | | ENSG00000272711 | AC019069.1 |
| | ENSG00000273237 | AC004520.1 | | ENSG00000266962 | AC067852.2 |
| | ENSG00000205485 | AC004980.1 | | ENSG00000101442 | ACTR5 |
| | ENSG00000224046 | AC005076.1 | | ENSG00000163485 | ADORA1 |
| | ENSG00000230882 | AC005077.4 | | ENSG00000258634 | AL160006.1 |
| | ENSG00000251537 | AC005324.3 | | ENSG00000151458 | ANKRD50 |
| | ENSG00000267056 | AC005336.1 | | ENSG00000122644 | ARL4A |
| | ENSG00000258653 | AC005520.1 | | ENSG00000114098 | ARMC8 |
| | ENSG00000261886 | AC005670.1 | | ENSG00000125962 | ARMCX5 |
| | ENSG00000266933 | AC005775.1 | | ENSG00000148219 | ASTN2 |
| | ENSG00000279452 | AC006277.1 | | ENSG00000119778 | ATAD2B |
| | ENSG00000277182 | AC006449.5 | | ENSG00000168646 | AXIN2 |
| | ENSG00000279539 | AC006486.2 | | ENSG00000174684 | B4GAT1 |
| | ENSG00000233937 | AC008443.1 | | ENSG00000125378 | BMP4 |
| | ENSG00000270067 | AC008536.1 | | ENSG00000215012 | C22orf29 |
| | ENSG00000197332 | AC008543.1 | | ENSG00000140326 | CDAN1 |
| | ENSG00000272869 | AC008575.2 | | ENSG00000165556 | CDX2 |
| | ENSG00000268655 | AC008687.4 | | ENSG00000147119 | CHST7 |
| | ENSG00000255441 | AC008750.2 | | ENSG00000198894 | CIPC |
| | ENSG00000269399 | AC008764.6 | | ENSG00000140931 | CMTM3 |
| | ENSG00000279529 | AC008764.8 | | ENSG00000118260 | CREB1 |
| | ENSG00000270020 | AC009108.3 | | ENSG00000144579 | CTDSP1 |
| | ENSG00000257176 | AC009318.1 | | ENSG00000175215 | CTDSP2 |
| | ENSG00000236255 | AC009404.1 | | ENSG00000171604 | CXXC5 |
| | ENSG00000269439 | AC010618.3 | | ENSG00000153071 | DAB2 |
| | ENSG00000267598 | AC011446.2 | | ENSG00000145358 | DDIT4L |
| | ENSG00000268583 | AC011466.1 | | ENSG00000116138 | DNAJC16 |
| | ENSG00000261777 | AC012184.3 | | ENSG00000110042 | DTX4 |
| | ENSG00000272994 | AC012360.3 | | ENSG00000108861 | DUSP3 |
| | ENSG00000260103 | AC012435.1 | | ENSG00000127334 | DYRK2 |
| | ENSG00000261526 | AC012615.1 | | ENSG00000151617 | EDNRA |
| | ENSG00000259802 | AC012640.2 | | ENSG00000243364 | EFNA4 |
| | ENSG00000250616 | AC012645.1 | | ENSG00000171617 | ENC1 |
| | ENSG00000258461 | AC012651.1 | | ENSG00000163508 | EOMES |
| | ENSG00000270195 | AC016773.1 | | ENSG00000187266 | EPOR |
| | ENSG00000273064 | AC017083.2 | | ENSG00000145569 | FAM105A |
| | ENSG00000273654 | AC020904.2 | | ENSG00000100376 | FAM118A |
| | ENSG00000283515 | AC020915.6 | | ENSG00000154319 | FAM167A |
| | ENSG00000267169 | AC022098.1 | | ENSG00000177706 | FAM20C |
| | ENSG00000260276 | AC022167.2 | | ENSG00000185614 | FAM212A |
| | ENSG00000279088 | AC022400.7 | | ENSG00000197852 | FAM212B |
| | ENSG00000259172 | AC023024.1 | | ENSG00000154511 | FAM69A |
| | ENSG00000257390 | AC023055.1 | | ENSG00000126882 | FAM78A |
| | ENSG00000223722 | AC023157.1 | | ENSG00000179431 | FJX1 |
| | ENSG00000261136 | AC023908.3 | | ENSG00000065970 | FOXJ2 |
| | ENSG00000236833 | AC024560.1 | | ENSG00000172728 | FUT10 |
| | ENSG00000272899 | AC025594.3 | | ENSG00000157240 | FZD1 |
| | ENSG00000260267 | AC026471.1 | | ENSG00000187210 | GCNT1 |
| | ENSG00000257894 | AC027288.3 | | ENSG00000142252 | GEMIN7 |
| | ENSG00000271918 | AC034236.2 | | ENSG00000189060 | H1F0 |
| | ENSG00000253200 | AC037459.3 | | ENSG00000103044 | HAS3 |
| | ENSG00000236814 | AC046176.1 | | ENSG00000135245 | HILPDA |
| | ENSG00000247373 | AC055713.1 | | ENSG00000275410 | HNF1B |

| | | | |
|---|---|---|---|
| ENSG00000234028 | AC062029.1 | ENSG00000166189 | HPS6 |
| ENSG00000240882 | AC063952.2 | ENSG00000002587 | HS3ST1 |
| ENSG00000274828 | AC068473.5 | ENSG00000115738 | ID2 |
| ENSG00000250644 | AC068580.4 | ENSG00000259330 | INAFM2 |
| ENSG00000214432 | AC068831.1 | ENSG00000185085 | INTS5 |
| ENSG00000272693 | AC073107.1 | ENSG00000168310 | IRF2 |
| ENSG00000273151 | AC073957.3 | ENSG00000165898 | ISCA2 |
| ENSG00000273391 | AC083880.1 | ENSG00000161638 | ITGA5 |
| ENSG00000274292 | AC084018.2 | ENSG00000198885 | ITPRIPL1 |
| ENSG00000257698 | AC084033.3 | ENSG00000151364 | KCTD14 |
| ENSG00000257732 | AC089983.1 | ENSG00000245060 | LINC00847 |
| ENSG00000272486 | AC090922.1 | ENSG00000136141 | LRCH1 |
| ENSG00000251127 | AC091173.1 | ENSG00000128011 | LRFN1 |
| ENSG00000273702 | AC091271.1 | ENSG00000185090 | MANEAL |
| ENSG00000229043 | AC091729.3 | ENSG00000095015 | MAP3K1 |
| ENSG00000198211 | AC092143.1 | ENSG00000112062 | MAPK14 |
| ENSG00000272275 | AC092687.3 | ENSG00000139266 | MARCH9 |
| ENSG00000260805 | AC092803.2 | ENSG00000277443 | MARCKS |
| ENSG00000246526 | AC093323.2 | ENSG00000247626 | MARS2 |
| ENSG00000226856 | AC093901.1 | ENSG00000124641 | MED20 |
| ENSG00000250673 | AC097372.1 | ENSG00000186260 | MKL2 |
| ENSG00000236432 | AC097662.1 | ENSG00000136997 | MYC |
| ENSG00000251615 | AC104825.3 | ENSG00000161653 | NAGS |
| ENSG00000260219 | AC106782.2 | ENSG00000074356 | NCBP3 |
| ENSG00000253741 | AC108002.1 | ENSG00000196865 | NHLRC2 |
| ENSG00000261889 | AC108134.2 | ENSG00000229544 | NKX1-2 |
| ENSG00000255224 | AC109322.1 | ENSG00000175745 | NR2F1 |
| ENSG00000275966 | AC110285.6 | ENSG00000185551 | NR2F2 |
| ENSG00000231305 | AC112484.2 | ENSG00000143867 | OSR1 |
| ENSG00000261159 | AC112484.4 | ENSG00000198754 | OXCT2 |
| ENSG00000263826 | AC112907.3 | ENSG00000175591 | P2RY2 |
| ENSG00000261351 | AC116913.1 | ENSG00000100105 | PATZ1 |
| ENSG00000279641 | AC120057.4 | ENSG00000243232 | PCDHAC2 |
| ENSG00000257497 | AC121761.1 | ENSG00000138735 | PDE5A |
| ENSG00000272677 | AC124016.1 | ENSG00000164951 | PDP1 |
| ENSG00000261441 | AC124068.2 | ENSG00000166821 | PEX11A |
| ENSG00000261924 | AC127496.1 | ENSG00000131779 | PEX11B |
| ENSG00000256092 | AC137767.1 | ENSG00000137338 | PGBD1 |
| ENSG00000278000 | AC139100.2 | ENSG00000174307 | PHLDA3 |
| ENSG00000260735 | AC139256.1 | ENSG00000274602 | PI4KAP1 |
| ENSG00000249592 | AC139887.2 | ENSG00000225973 | PIGBOS1 |
| ENSG00000230650 | AC140479.2 | ENSG00000120278 | PLEKHG1 |
| ENSG00000276728 | AC142472.1 | ENSG00000126822 | PLEKHG3 |
| ENSG00000261888 | AC144831.1 | ENSG00000171453 | POLR1C |
| ENSG00000270012 | AC232271.1 | ENSG00000111110 | PPM1H |
| ENSG00000261716 | AC239868.2 | ENSG00000108819 | PPP1R9B |
| ENSG00000264207 | AC239868.3 | ENSG00000275342 | PRAG1 |
| ENSG00000233586 | AC246785.2 | ENSG00000237943 | PRKCQ-AS1 |
| ENSG00000111644 | ACRBP | ENSG00000198890 | PRMT6 |
| ENSG00000130377 | ACSBG2 | ENSG00000116574 | RHOU |
| ENSG00000143632 | ACTA1 | ENSG00000107036 | RIC1 |
| ENSG00000107796 | ACTA2 | ENSG00000108375 | RNF43 |
| ENSG00000178631 | ACTG1P1 | ENSG00000169071 | ROR2 |
| ENSG00000241547 | ACTG1P20 | ENSG00000179041 | RRS1 |
| ENSG00000117148 | ACTL8 | ENSG00000185924 | RTN4RL1 |
| ENSG00000158859 | ADAMTS4 | ENSG00000198863 | RUNDC1 |
| ENSG00000143382 | ADAMTSL4 | ENSG00000180739 | S1PR5 |
| ENSG00000161912 | ADCY10P1 | ENSG00000185033 | SEMA4B |
| ENSG00000111452 | ADGRD1 | ENSG00000104611 | SH2D4A |
| ENSG00000147576 | ADHFE1 | ENSG00000145248 | SLC10A4 |

| | |
|---|---|
| ENSG00000153531 | ADPRHL1 |
| ENSG00000170222 | ADPRM |
| ENSG00000274286 | ADRA2B |
| ENSG00000280273 | AF131216.4 |
| ENSG00000172650 | AGAP5 |
| ENSG00000146856 | AGBL3 |
| ENSG00000197976 | AKAP17A |
| ENSG00000179841 | AKAP5 |
| ENSG00000135334 | AKIRIN2 |
| ENSG00000284431 | AL022238.4 |
| ENSG00000226954 | AL023802.1 |
| ENSG00000272277 | AL031963.3 |
| ENSG00000230424 | AL035413.1 |
| ENSG00000279267 | AL078621.3 |
| ENSG00000227627 | AL080276.2 |
| ENSG00000268858 | AL118506.1 |
| ENSG00000260708 | AL118516.1 |
| ENSG00000279253 | AL121753.2 |
| ENSG00000273619 | AL121832.2 |
| ENSG00000271361 | AL133351.4 |
| ENSG00000258727 | AL135999.1 |
| ENSG00000241255 | AL136126.1 |
| ENSG00000283078 | AL137077.2 |
| ENSG00000261135 | AL137802.2 |
| ENSG00000260193 | AL138781.1 |
| ENSG00000280710 | AL139035.1 |
| ENSG00000273723 | AL139089.1 |
| ENSG00000187186 | AL162231.1 |
| ENSG00000261215 | AL162231.4 |
| ENSG00000260806 | AL163051.1 |
| ENSG00000223503 | AL353807.1 |
| ENSG00000236345 | AL354719.2 |
| ENSG00000254473 | AL354920.1 |
| ENSG00000205740 | AL359878.1 |
| ENSG00000273058 | AL359921.2 |
| ENSG00000272068 | AL365181.2 |
| ENSG00000229656 | AL365203.1 |
| ENSG00000272455 | AL391244.3 |
| ENSG00000224934 | AL391684.1 |
| ENSG00000276248 | AL442125.1 |
| ENSG00000240291 | AL450384.2 |
| ENSG00000228192 | AL512353.1 |
| ENSG00000256966 | AL513165.2 |
| ENSG00000279443 | AL513497.1 |
| ENSG00000227603 | AL583839.1 |
| ENSG00000238009 | AL627309.1 |
| ENSG00000223764 | AL645608.1 |
| ENSG00000272217 | AL645940.1 |
| ENSG00000280128 | AL662795.2 |
| ENSG00000237491 | AL669831.5 |
| ENSG00000272106 | AL691432.2 |
| ENSG00000273599 | AL731571.1 |
| ENSG00000227518 | AL928970.1 |
| ENSG00000163631 | ALB |
| ENSG00000240038 | AMY2B |
| ENSG00000261239 | ANKRD26P1 |
| ENSG00000145700 | ANKRD31 |
| ENSG00000166825 | ANPEP |
| ENSG00000131471 | AOC3 |
| ENSG00000138356 | AOX1 |
| ENSG00000115840 | SLC25A12 |
| ENSG00000198246 | SLC29A3 |
| ENSG00000170385 | SLC30A1 |
| ENSG00000100036 | SLC35E4 |
| ENSG00000170365 | SMAD1 |
| ENSG00000124104 | SNX21 |
| ENSG00000173548 | SNX33 |
| ENSG00000124766 | SOX4 |
| ENSG00000125398 | SOX9 |
| ENSG00000166068 | SPRED1 |
| ENSG00000136158 | SPRY2 |
| ENSG00000143093 | STRIP1 |
| ENSG00000139531 | SUOX |
| ENSG00000100324 | TAB1 |
| ENSG00000168769 | TET2 |
| ENSG00000130193 | THEM6 |
| ENSG00000177370 | TIMM22 |
| ENSG00000144120 | TMEM177 |
| ENSG00000165152 | TMEM246 |
| ENSG00000149115 | TNKS1BP1 |
| ENSG00000146242 | TPBG |
| ENSG00000204977 | TRIM13 |
| ENSG00000116525 | TRIM62 |
| ENSG00000102804 | TSC22D1 |
| ENSG00000182704 | TSKU |
| ENSG00000182040 | USH1G |
| ENSG00000273820 | USP27X |
| ENSG00000176428 | VPS37D |
| ENSG00000139131 | YARS2 |
| ENSG00000180011 | ZADH2 |
| ENSG00000177494 | ZBED2 |
| ENSG00000213762 | ZNF134 |
| ENSG00000185219 | ZNF445 |
| ENSG00000265763 | ZNF488 |
| ENSG00000162714 | ZNF496 |
| ENSG00000218891 | ZNF579 |
| ENSG00000083828 | ZNF586 |
| ENSG00000102870 | ZNF629 |
| ENSG00000198093 | ZNF649 |
| ENSG00000196453 | ZNF777 |
| ENSG00000196812 | ZSCAN16 |

| | |
|---|---|
| ENSG00000284128 | AP000356.3 |
| ENSG00000280594 | AP000432.3 |
| ENSG00000254979 | AP000781.2 |
| ENSG00000247137 | AP000873.2 |
| ENSG00000269176 | AP001160.3 |
| ENSG00000256116 | AP001453.1 |
| ENSG00000280367 | AP002364.1 |
| ENSG00000266401 | AP002478.1 |
| ENSG00000255508 | AP002990.1 |
| ENSG00000253669 | AP003356.1 |
| ENSG00000255553 | AP003733.3 |
| ENSG00000115266 | APC2 |
| ENSG00000130203 | APOE |
| ENSG00000221963 | APOL6 |
| ENSG00000204444 | APOM |
| ENSG00000240583 | AQP1 |
| ENSG00000111348 | ARHGDIB |
| ENSG00000183111 | ARHGEF37 |
| ENSG00000221883 | ARIH2OS |
| ENSG00000179674 | ARL14 |
| ENSG00000268714 | ARL14EPL |
| ENSG00000196503 | ARL9 |
| ENSG00000203993 | ARRDC1-AS1 |
| ENSG00000113369 | ARRDC3 |
| ENSG00000141505 | ASGR1 |
| ENSG00000072182 | ASIC4 |
| ENSG00000244617 | ASPRV1 |
| ENSG00000188886 | ASTL |
| ENSG00000162772 | ATF3 |
| ENSG00000213760 | ATP6V1G2 |
| ENSG00000253320 | AZIN1-AS1 |
| ENSG00000247081 | BAALC-AS1 |
| ENSG00000136881 | BAAT |
| ENSG00000151929 | BAG3 |
| ENSG00000131668 | BARX1 |
| ENSG00000245573 | BDNF-AS |
| ENSG00000237452 | BHMG1 |
| ENSG00000010671 | BTK |
| ENSG00000186265 | BTLA |
| ENSG00000260996 | BX255925.1 |
| ENSG00000180425 | C11orf71 |
| ENSG00000214226 | C17orf67 |
| ENSG00000187997 | C17orf99 |
| ENSG00000105072 | C19orf44 |
| ENSG00000262874 | C19orf84 |
| ENSG00000145861 | C1QTNF2 |
| ENSG00000182326 | C1S |
| ENSG00000166278 | C2 |
| ENSG00000196421 | C20orf204 |
| ENSG00000188511 | C22orf34 |
| ENSG00000115998 | C2orf42 |
| ENSG00000172478 | C2orf54 |
| ENSG00000187833 | C2orf78 |
| ENSG00000125730 | C3 |
| ENSG00000244731 | C4A |
| ENSG00000224389 | C4B |
| ENSG00000197405 | C5AR1 |
| ENSG00000189325 | C6orf222 |
| ENSG00000270024 | C8orf44-SGK3 |
| ENSG00000169085 | C8orf46 |

| | |
|---|---|
| ENSG00000196366 | C9orf163 |
| ENSG00000186312 | CA5BP1 |
| ENSG00000260942 | CAPN10-AS1 |
| ENSG00000092529 | CAPN3 |
| ENSG00000180881 | CAPS2 |
| ENSG00000187796 | CARD9 |
| ENSG00000118412 | CASP8AP2 |
| ENSG00000087589 | CASS4 |
| ENSG00000099625 | CBARP |
| ENSG00000204659 | CBY3 |
| ENSG00000173581 | CCDC106 |
| ENSG00000256304 | CCDC150P1 |
| ENSG00000198003 | CCDC151 |
| ENSG00000130783 | CCDC62 |
| ENSG00000119242 | CCDC92 |
| ENSG00000132141 | CCT6B |
| ENSG00000170458 | CD14 |
| ENSG00000163606 | CD200R1 |
| ENSG00000129226 | CD68 |
| ENSG00000125726 | CD70 |
| ENSG00000167797 | CDK2AP2 |
| ENSG00000006837 | CDKL3 |
| ENSG00000129355 | CDKN2D |
| ENSG00000100526 | CDKN3 |
| ENSG00000129596 | CDO1 |
| ENSG00000241322 | CDRT1 |
| ENSG00000277449 | CEBPB-AS1 |
| ENSG00000008300 | CELSR3 |
| ENSG00000184524 | CEND1 |
| ENSG00000177946 | CENPBD1 |
| ENSG00000087237 | CETP |
| ENSG00000197748 | CFAP43 |
| ENSG00000172361 | CFAP53 |
| ENSG00000120051 | CFAP58 |
| ENSG00000181378 | CFAP65 |
| ENSG00000243649 | CFB |
| ENSG00000120903 | CHRNA2 |
| ENSG00000101204 | CHRNA4 |
| ENSG00000186162 | CIDECP |
| ENSG00000267493 | CIRBP-AS1 |
| ENSG00000175505 | CLCF1 |
| ENSG00000184697 | CLDN6 |
| ENSG00000013441 | CLK1 |
| ENSG00000140932 | CMTM2 |
| ENSG00000149646 | CNBD2 |
| ENSG00000143786 | CNIH3 |
| ENSG00000136152 | COG3 |
| ENSG00000101203 | COL20A1 |
| ENSG00000139219 | COL2A1 |
| ENSG00000145244 | CORIN |
| ENSG00000134376 | CRB1 |
| ENSG00000143578 | CREB3L4 |
| ENSG00000103196 | CRISPLD2 |
| ENSG00000268916 | CSAG3 |
| ENSG00000114646 | CSPG5 |
| ENSG00000118523 | CTGF |
| ENSG00000280018 | CU634019.6 |
| ENSG00000280145 | CU638689.4 |
| ENSG00000138161 | CUZD1 |
| ENSG00000168329 | CX3CR1 |

| | |
|---|---|
| ENSG00000161544 | CYGB |
| ENSG00000137869 | CYP19A1 |
| ENSG00000135929 | CYP27A1 |
| ENSG00000256612 | CYP2B7P |
| ENSG00000171903 | CYP4F11 |
| ENSG00000142871 | CYR61 |
| ENSG00000205795 | CYS1 |
| ENSG00000197191 | CYSRT1 |
| ENSG00000182308 | DCAF4L1 |
| ENSG00000166341 | DCHS1 |
| ENSG00000175197 | DDIT3 |
| ENSG00000162733 | DDR2 |
| ENSG00000109832 | DDX25 |
| ENSG00000160570 | DEDD2 |
| ENSG00000197406 | DIO3 |
| ENSG00000268471 | DKFZP434I0714 |
| ENSG00000132535 | DLG4 |
| ENSG00000170579 | DLGAP1 |
| ENSG00000132837 | DMGDH |
| ENSG00000154099 | DNAAF1 |
| ENSG00000114841 | DNAH1 |
| ENSG00000197653 | DNAH10 |
| ENSG00000174844 | DNAH12 |
| ENSG00000187775 | DNAH17 |
| ENSG00000086061 | DNAJA1 |
| ENSG00000140403 | DNAJA4 |
| ENSG00000132002 | DNAJB1 |
| ENSG00000247400 | DNAJC3-AS1 |
| ENSG00000179532 | DNHD1 |
| ENSG00000157856 | DRC1 |
| ENSG00000159625 | DRC7 |
| ENSG00000102385 | DRP2 |
| ENSG00000158050 | DUSP2 |
| ENSG00000188542 | DUSP28 |
| ENSG00000184545 | DUSP8 |
| ENSG00000235316 | DUSP8P5 |
| ENSG00000172771 | EFCAB12 |
| ENSG00000172638 | EFEMP2 |
| ENSG00000115468 | EFHD1 |
| ENSG00000120738 | EGR1 |
| ENSG00000122877 | EGR2 |
| ENSG00000179388 | EGR3 |
| ENSG00000135625 | EGR4 |
| ENSG00000255150 | EID3 |
| ENSG00000172071 | EIF2AK3 |
| ENSG00000196361 | ELAVL3 |
| ENSG00000180385 | EMC3-AS1 |
| ENSG00000108515 | ENO3 |
| ENSG00000136960 | ENPP2 |
| ENSG00000116106 | EPHA4 |
| ENSG00000164010 | ERMAP |
| ENSG00000260565 | ERVK13-1 |
| ENSG00000144488 | ESPNL |
| ENSG00000119715 | ESRRB |
| ENSG00000187609 | EXD3 |
| ENSG00000221990 | EXOC3-AS1 |
| ENSG00000057593 | F7 |
| ENSG00000131944 | FAAP24 |
| ENSG00000121104 | FAM117A |
| ENSG00000184083 | FAM120C |

| | |
|---|---|
| ENSG00000047662 | FAM184B |
| ENSG00000219626 | FAM228B |
| ENSG00000283709 | FAM238C |
| ENSG00000174137 | FAM53A |
| ENSG00000149926 | FAM57B |
| ENSG00000125998 | FAM83C |
| ENSG00000188573 | FBLL1 |
| ENSG00000147912 | FBXO10 |
| ENSG00000141665 | FBXO15 |
| ENSG00000156509 | FBXO43 |
| ENSG00000171931 | FBXW10 |
| ENSG00000275395 | FCGBP |
| ENSG00000162746 | FCRLB |
| ENSG00000214814 | FER1L6 |
| ENSG00000149781 | FERMT3 |
| ENSG00000149557 | FEZ1 |
| ENSG00000142621 | FHAD1 |
| ENSG00000198855 | FICD |
| ENSG00000224023 | FLJ37035 |
| ENSG00000179743 | FLJ37453 |
| ENSG00000248905 | FMN1 |
| ENSG00000076258 | FMO4 |
| ENSG00000272004 | FO704657.1 |
| ENSG00000170345 | FOS |
| ENSG00000125740 | FOSB |
| ENSG00000187559 | FOXD4L3 |
| ENSG00000184659 | FOXD4L4 |
| ENSG00000049768 | FOXP3 |
| ENSG00000279177 | FP236241.2 |
| ENSG00000172159 | FRMD3 |
| ENSG00000154727 | GABPA |
| ENSG00000130222 | GADD45G |
| ENSG00000182687 | GALR2 |
| ENSG00000160766 | GBAP1 |
| ENSG00000137270 | GCM1 |
| ENSG00000147174 | GCNA |
| ENSG00000164404 | GDF9 |
| ENSG00000228175 | GEMIN8P4 |
| ENSG00000100031 | GGT1 |
| ENSG00000162419 | GMEB1 |
| ENSG00000173540 | GMPPB |
| ENSG00000159289 | GOLGA6A |
| ENSG00000136235 | GPNMB |
| ENSG00000277399 | GPR179 |
| ENSG00000183150 | GPR19 |
| ENSG00000178623 | GPR35 |
| ENSG00000123901 | GPR83 |
| ENSG00000023171 | GRAMD1B |
| ENSG00000154016 | GRAP |
| ENSG00000105737 | GRIK5 |
| ENSG00000161509 | GRIN2C |
| ENSG00000196275 | GTF2IRD2 |
| ENSG00000206172 | HBA1 |
| ENSG00000188536 | HBA2 |
| ENSG00000196565 | HBG2 |
| ENSG00000099822 | HCN2 |
| ENSG00000051108 | HERPUD1 |
| ENSG00000164683 | HEY1 |
| ENSG00000177374 | HIC1 |
| ENSG00000204257 | HLA-DMA |

| | |
|---|---|
| ENSG00000242574 | HLA-DMB |
| ENSG00000219163 | HMGB1P20 |
| ENSG00000103942 | HOMER2 |
| ENSG00000233429 | HOTAIRM1 |
| ENSG00000158104 | HPD |
| ENSG00000110169 | HPX |
| ENSG00000113905 | HRG |
| ENSG00000025423 | HSD17B6 |
| ENSG00000204389 | HSPA1A |
| ENSG00000204388 | HSPA1B |
| ENSG00000204390 | HSPA1L |
| ENSG00000173110 | HSPA6 |
| ENSG00000260325 | HSPB9 |
| ENSG00000120694 | HSPH1 |
| ENSG00000067064 | IDI1 |
| ENSG00000160888 | IER2 |
| ENSG00000099869 | IGF2-AS |
| ENSG00000211895 | IGHA1 |
| ENSG00000211890 | IGHA2 |
| ENSG00000211891 | IGHE |
| ENSG00000211893 | IGHG2 |
| ENSG00000095752 | IL11 |
| ENSG00000168811 | IL12A |
| ENSG00000172349 | IL16 |
| ENSG00000112116 | IL17F |
| ENSG00000110944 | IL23A |
| ENSG00000157368 | IL34 |
| ENSG00000113525 | IL5 |
| ENSG00000232133 | IMPDH1P10 |
| ENSG00000153487 | ING1 |
| ENSG00000175189 | INHBC |
| ENSG00000186480 | INSIG1 |
| ENSG00000236778 | INTS6-AS1 |
| ENSG00000174628 | IQCK |
| ENSG00000090376 | IRAK3 |
| ENSG00000100593 | ISM2 |
| ENSG00000213949 | ITGA1 |
| ENSG00000169896 | ITGAM |
| ENSG00000154118 | JPH3 |
| ENSG00000177606 | JUN |
| ENSG00000171223 | JUNB |
| ENSG00000167216 | KATNAL2 |
| ENSG00000107821 | KAZALD1 |
| ENSG00000123444 | KBTBD4 |
| ENSG00000162728 | KCNJ9 |
| ENSG00000171303 | KCNK3 |
| ENSG00000105642 | KCNN1 |
| ENSG00000213859 | KCTD11 |
| ENSG00000183775 | KCTD16 |
| ENSG00000246174 | KCTD21-AS1 |
| ENSG00000128052 | KDR |
| ENSG00000196872 | KIAA1211L |
| ENSG00000130518 | KIAA1683 |
| ENSG00000226650 | KIF4B |
| ENSG00000155090 | KLF10 |
| ENSG00000124743 | KLHL31 |
| ENSG00000164344 | KLKB1 |
| ENSG00000186395 | KRT10 |
| ENSG00000171401 | KRT13 |
| ENSG00000205420 | KRT6A |

| | |
|---|---|
| ENSG00000229320 | KRT8P12 |
| ENSG00000115850 | LCT |
| ENSG00000168675 | LDLRAD4 |
| ENSG00000164406 | LEAP2 |
| ENSG00000197980 | LEKR1 |
| ENSG00000107187 | LHX3 |
| ENSG00000225760 | LINC00431 |
| ENSG00000229373 | LINC00452 |
| ENSG00000233559 | LINC00513 |
| ENSG00000179935 | LINC00652 |
| ENSG00000228705 | LINC00659 |
| ENSG00000231711 | LINC00899 |
| ENSG00000240476 | LINC00973 |
| ENSG00000228393 | LINC01004 |
| ENSG00000245146 | LINC01024 |
| ENSG00000212694 | LINC01089 |
| ENSG00000239332 | LINC01119 |
| ENSG00000280734 | LINC01232 |
| ENSG00000203999 | LINC01270 |
| ENSG00000260924 | LINC01311 |
| ENSG00000183250 | LINC01547 |
| ENSG00000262468 | LINC01569 |
| ENSG00000233396 | LINC01719 |
| ENSG00000262155 | LINC02175 |
| ENSG00000271538 | LINC02427 |
| ENSG00000231721 | LINC-PINT |
| ENSG00000136153 | LMO7 |
| ENSG00000163431 | LMOD1 |
| ENSG00000206535 | LNP1 |
| ENSG00000139679 | LPAR6 |
| ENSG00000183423 | LRIT3 |
| ENSG00000109771 | LRP2BP |
| ENSG00000247675 | LRP4-AS1 |
| ENSG00000176809 | LRRC37A3 |
| ENSG00000188993 | LRRC66 |
| ENSG00000226979 | LTA |
| ENSG00000119681 | LTBP2 |
| ENSG00000198862 | LTN1 |
| ENSG00000160886 | LY6K |
| ENSG00000198798 | MAGEB3 |
| ENSG00000006062 | MAP3K14 |
| ENSG00000186056 | MATN1-AS1 |
| ENSG00000258839 | MC1R |
| ENSG00000102802 | MEDAG |
| ENSG00000197889 | MEIG1 |
| ENSG00000092931 | MFSD11 |
| ENSG00000257335 | MGAM |
| ENSG00000186594 | MIR22HG |
| ENSG00000260083 | MIR762HG |
| ENSG00000167842 | MIS12 |
| ENSG00000076242 | MLH1 |
| ENSG00000008516 | MMP25 |
| ENSG00000168314 | MOBP |
| ENSG00000228782 | MRPL45P2 |
| ENSG00000138379 | MSTN |
| ENSG00000198804 | MT-CO1 |
| ENSG00000120662 | MTRF1 |
| ENSG00000138823 | MTTP |
| ENSG00000179141 | MTUS2-AS1 |
| ENSG00000176945 | MUC20 |

| | |
|---|---|
| ENSG00000224769 | MUC20P1 |
| ENSG00000215182 | MUC5AC |
| ENSG00000059728 | MXD1 |
| ENSG00000162576 | MXRA8 |
| ENSG00000185105 | MYADML2 |
| ENSG00000109063 | MYH3 |
| ENSG00000101306 | MYLK2 |
| ENSG00000166866 | MYO1A |
| ENSG00000036448 | MYOM2 |
| ENSG00000138347 | MYPN |
| ENSG00000099326 | MZF1 |
| ENSG00000267858 | MZF1-AS1 |
| ENSG00000136274 | NACAD |
| ENSG00000171169 | NAIF1 |
| ENSG00000067798 | NAV3 |
| ENSG00000272150 | NBPF25P |
| ENSG00000130287 | NCAN |
| ENSG00000183091 | NEB |
| ENSG00000114670 | NEK11 |
| ENSG00000230257 | NFE4 |
| ENSG00000144802 | NFKBIZ |
| ENSG00000272145 | NFYC-AS1 |
| ENSG00000171786 | NHLH1 |
| ENSG00000179846 | NKPD1 |
| ENSG00000091106 | NLRC4 |
| ENSG00000160505 | NLRP4 |
| ENSG00000167634 | NLRP7 |
| ENSG00000015520 | NPC1L1 |
| ENSG00000254206 | NPIPB11 |
| ENSG00000198156 | NPIPB6 |
| ENSG00000233232 | NPIPB7 |
| ENSG00000196993 | NPIPB9 |
| ENSG00000123358 | NR4A1 |
| ENSG00000153234 | NR4A2 |
| ENSG00000119508 | NR4A3 |
| ENSG00000197893 | NRAP |
| ENSG00000091129 | NRCAM |
| ENSG00000110076 | NRXN2 |
| ENSG00000226328 | NUP50-AS1 |
| ENSG00000188199 | NUTM2B |
| ENSG00000188039 | NWD1 |
| ENSG00000101888 | NXT2 |
| ENSG00000177989 | ODF3B |
| ENSG00000235823 | OLMALINC |
| ENSG00000235213 | OR6E1P |
| ENSG00000223891 | OSER1-AS1 |
| ENSG00000085465 | OVGP1 |
| ENSG00000172818 | OVOL1 |
| ENSG00000159339 | PADI4 |
| ENSG00000135473 | PAN2 |
| ENSG00000274897 | PANO1 |
| ENSG00000267270 | PARD6G-AS1 |
| ENSG00000178685 | PARP10 |
| ENSG00000204956 | PCDHGA1 |
| ENSG00000102109 | PCSK1N |
| ENSG00000105650 | PDE4C |
| ENSG00000185527 | PDE6G |
| ENSG00000182405 | PGBD4 |
| ENSG00000119630 | PGF |
| ENSG00000161031 | PGLYRP2 |

| | |
|---|---|
| ENSG00000139200 | PIANP |
| ENSG00000085514 | PILRA |
| ENSG00000137193 | PIM1 |
| ENSG00000197181 | PIWIL2 |
| ENSG00000158683 | PKD1L1 |
| ENSG00000130943 | PKDREJ |
| ENSG00000105499 | PLA2G4C |
| ENSG00000184381 | PLA2G6 |
| ENSG00000115956 | PLEK |
| ENSG00000175985 | PLEKHD1 |
| ENSG00000100979 | PLTP |
| ENSG00000146453 | PNLDC1 |
| ENSG00000130653 | PNPLA7 |
| ENSG00000121577 | POPDC2 |
| ENSG00000028277 | POU2F2 |
| ENSG00000204569 | PPP1R10 |
| ENSG00000257557 | PPP1R12A-AS1 |
| ENSG00000087074 | PPP1R15A |
| ENSG00000182676 | PPP1R27 |
| ENSG00000049769 | PPP1R3F |
| ENSG00000230510 | PPP5D1 |
| ENSG00000126856 | PRDM7 |
| ENSG00000126460 | PRRG2 |
| ENSG00000172460 | PRSS30P |
| ENSG00000167653 | PSCA |
| ENSG00000169403 | PTAFR |
| ENSG00000050628 | PTGER3 |
| ENSG00000073756 | PTGS2 |
| ENSG00000163661 | PTX3 |
| ENSG00000172733 | PURG |
| ENSG00000179165 | PXT1 |
| ENSG00000068976 | PYGM |
| ENSG00000109113 | RAB34 |
| ENSG00000245849 | RAD51-AS1 |
| ENSG00000228265 | RALY-AS1 |
| ENSG00000197291 | RAMP2-AS1 |
| ENSG00000131759 | RARA |
| ENSG00000108551 | RASD1 |
| ENSG00000100302 | RASD2 |
| ENSG00000058335 | RASGRF1 |
| ENSG00000270885 | RASL10B |
| ENSG00000162775 | RBM15 |
| ENSG00000170748 | RBMXL2 |
| ENSG00000163918 | RFC4 |
| ENSG00000128253 | RFPL2 |
| ENSG00000128408 | RIBC2 |
| ENSG00000245149 | RNF139-AS1 |
| ENSG00000233198 | RNF224 |
| ENSG00000245534 | RORA-AS1 |
| ENSG00000215472 | RPL17-C18orf32 |
| ENSG00000140986 | RPL3L |
| ENSG00000241749 | RPSAP52 |
| ENSG00000166592 | RRAD |
| ENSG00000143303 | RRNAD1 |
| ENSG00000155026 | RSPH10B |
| ENSG00000163993 | S100P |
| ENSG00000187634 | SAMD11 |
| ENSG00000004139 | SARM1 |
| ENSG00000188659 | SAXO2 |
| ENSG00000231274 | SBK3 |

| | |
|---|---|
| ENSG00000189001 | SBSN |
| ENSG00000047634 | SCML1 |
| ENSG00000163156 | SCNM1 |
| ENSG00000284194 | SCO2 |
| ENSG00000175356 | SCUBE2 |
| ENSG00000234684 | SDCBP2-AS1 |
| ENSG00000143416 | SELENBP1 |
| ENSG00000110876 | SELPLG |
| ENSG00000184716 | SERINC4 |
| ENSG00000106366 | SERPINE1 |
| ENSG00000104897 | SF3A2 |
| ENSG00000148082 | SHC3 |
| ENSG00000185634 | SHC4 |
| ENSG00000187902 | SHISA7 |
| ENSG00000108932 | SLC16A6 |
| ENSG00000218363 | SLC25A20P1 |
| ENSG00000269743 | SLC25A53 |
| ENSG00000109667 | SLC2A9 |
| ENSG00000198569 | SLC34A3 |
| ENSG00000111181 | SLC6A12 |
| ENSG00000010379 | SLC6A13 |
| ENSG00000155465 | SLC7A7 |
| ENSG00000021488 | SLC7A9 |
| ENSG00000118160 | SLC8A2 |
| ENSG00000135740 | SLC9A5 |
| ENSG00000254634 | SMG1P6 |
| ENSG00000261556 | SMG1P7 |
| ENSG00000163683 | SMIM14 |
| ENSG00000019549 | SNAI2 |
| ENSG00000234912 | SNHG20 |
| ENSG00000250988 | SNHG21 |
| ENSG00000163877 | SNIP1 |
| ENSG00000039600 | SOX30 |
| ENSG00000167182 | SP2 |
| ENSG00000163806 | SPDYA |
| ENSG00000063176 | SPHK2 |
| ENSG00000107742 | SPOCK2 |
| ENSG00000118785 | SPP1 |
| ENSG00000188766 | SPRED3 |
| ENSG00000196220 | SRGAP3 |
| ENSG00000248508 | SRP14-AS1 |
| ENSG00000205913 | SRRM2-AS1 |
| ENSG00000139767 | SRRM4 |
| ENSG00000226763 | SRRM5 |
| ENSG00000260233 | SSSCA1-AS1 |
| ENSG00000125046 | SSUH2 |
| ENSG00000126091 | ST3GAL3 |
| ENSG00000178078 | STAP2 |
| ENSG00000168439 | STIP1 |
| ENSG00000248278 | SUMO2P17 |
| ENSG00000159164 | SV2A |
| ENSG00000105467 | SYNGR4 |
| ENSG00000166317 | SYNPO2L |
| ENSG00000102003 | SYP |
| ENSG00000132718 | SYT11 |
| ENSG00000173227 | SYT12 |
| ENSG00000213023 | SYT3 |
| ENSG00000102387 | TAF7L |
| ENSG00000164691 | TAGAP |
| ENSG00000144834 | TAGLN3 |

| | |
|---|---|
| ENSG00000136535 | TBR1 |
| ENSG00000121075 | TBX4 |
| ENSG00000173991 | TCAP |
| ENSG00000166046 | TCP11L2 |
| ENSG00000184786 | TCTE3 |
| ENSG00000109927 | TECTA |
| ENSG00000120156 | TEK |
| ENSG00000092850 | TEKT2 |
| ENSG00000121101 | TEX14 |
| ENSG00000182459 | TEX19 |
| ENSG00000042832 | TG |
| ENSG00000173451 | THAP2 |
| ENSG00000168152 | THAP9 |
| ENSG00000137801 | THBS1 |
| ENSG00000113296 | THBS4 |
| ENSG00000206573 | THUMPD3-AS1 |
| ENSG00000173825 | TIGD3 |
| ENSG00000169989 | TIGD4 |
| ENSG00000095587 | TLL2 |
| ENSG00000271270 | TMCC1-AS1 |
| ENSG00000251201 | TMED7-TICAM2 |
| ENSG00000198270 | TMEM116 |
| ENSG00000183160 | TMEM119 |
| ENSG00000161558 | TMEM143 |
| ENSG00000247828 | TMEM161B-AS1 |
| ENSG00000172738 | TMEM217 |
| ENSG00000231770 | TMEM44-AS1 |
| ENSG00000163472 | TMEM79 |
| ENSG00000262904 | TMPOP2 |
| ENSG00000178297 | TMPRSS9 |
| ENSG00000028137 | TNFRSF1B |
| ENSG00000120949 | TNFRSF8 |
| ENSG00000161955 | TNFSF13 |
| ENSG00000168477 | TNXB |
| ENSG00000230359 | TPI1P2 |
| ENSG00000234127 | TRIM26 |
| ENSG00000248167 | TRIM39-RPP21 |
| ENSG00000132256 | TRIM5 |
| ENSG00000178809 | TRIM73 |
| ENSG00000089195 | TRMT6 |
| ENSG00000167723 | TRPV3 |
| ENSG00000163467 | TSACC |
| ENSG00000005379 | TSPOAP1 |
| ENSG00000178093 | TSSK6 |
| ENSG00000235954 | TTC28-AS1 |
| ENSG00000183891 | TTC32 |
| ENSG00000113638 | TTC33 |
| ENSG00000214198 | TTC41P |
| ENSG00000155657 | TTN |
| ENSG00000258947 | TUBB3 |
| ENSG00000261812 | TUBB8P7 |
| ENSG00000160201 | U2AF1 |
| ENSG00000272821 | U62317.3 |
| ENSG00000182179 | UBA7 |
| ENSG00000150991 | UBC |
| ENSG00000175518 | UBQLNL |
| ENSG00000104691 | UBXN8 |
| ENSG00000280213 | UCKL1-AS1 |
| ENSG00000175564 | UCP3 |
| ENSG00000249348 | UGDH-AS1 |

| | |
|---|---|
| ENSG00000124602 | UNC5CL |
| ENSG00000132952 | USPL1 |
| ENSG00000163945 | UVSSA |
| ENSG00000165197 | VEGFD |
| ENSG00000119614 | VSX2 |
| ENSG00000015285 | WAS |
| ENSG00000047056 | WDR37 |
| ENSG00000158023 | WDR66 |
| ENSG00000152763 | WDR78 |
| ENSG00000127578 | WFIKKN1 |
| ENSG00000196632 | WNK3 |
| ENSG00000100219 | XBP1 |
| ENSG00000196449 | YRDC |
| ENSG00000168826 | ZBTB49 |
| ENSG00000163874 | ZC3H12A |
| ENSG00000178381 | ZFAND2A |
| ENSG00000128016 | ZFP36 |
| ENSG00000166478 | ZNF143 |
| ENSG00000263072 | ZNF213-AS1 |
| ENSG00000159915 | ZNF233 |
| ENSG00000198169 | ZNF251 |
| ENSG00000160961 | ZNF333 |
| ENSG00000196378 | ZNF34 |
| ENSG00000144331 | ZNF385B |
| ENSG00000204947 | ZNF425 |
| ENSG00000249087 | ZNF436-AS1 |
| ENSG00000173258 | ZNF483 |
| ENSG00000188033 | ZNF490 |
| ENSG00000187187 | ZNF546 |
| ENSG00000152433 | ZNF547 |
| ENSG00000172748 | ZNF596 |
| ENSG00000197483 | ZNF628 |
| ENSG00000197054 | ZNF763 |
| ENSG00000152475 | ZNF837 |
| ENSG00000176723 | ZNF843 |
| ENSG00000196605 | ZNF846 |
| ENSG00000188372 | ZP3 |
| ENSG00000130182 | ZSCAN10 |
| ENSG00000219891 | ZSCAN12P1 |
| ENSG00000235109 | ZSCAN31 |
